# jazzPanda: spatially aware marker gene detection for imaging-based spatial transcriptomics

**DOI:** 10.64898/2026.02.13.705867

**Authors:** Xinyi Jin, Givanna H. Putri, Jinming Cheng, Marie-Liesse Asselin-Labat, Gordon Smyth, Belinda Phipson

## Abstract

**Motivation:** Spatial transcriptomics resolves the organisation of tissues and their cellular neighbourhoods, where cell type identification depends on reliable marker gene detection. Existing marker methods were developed for single-cell RNA sequencing and ignore the spatial coordinates of cells and transcripts, a particular problem for imaging-based platforms where transcript counts per gene per cell are extremely sparse.

**Results:** We present jazzPanda, a method for detecting spatially informative marker genes in imaging-based spatial transcriptomics. Transcript and cell coordinates are aggregated into one-dimensional vectors by spatial binning, and gene vectors can be built directly from transcript coordinates without cell segmentation. Markers are identified either by permutation-based rank correlation for single-sample data, or by a lasso-regularised generalised linear model that accommodates multiple samples and platform-specific background signal from negative-control probes. To our knowledge, jazzPanda is the only marker detection method to account jointly for spatial distribution, replication across samples, and platform background. Benchmarked against the Wilcoxon rank sum test and t-tests on public Xenium and CosMx data, jazzPanda recovers markers with stronger spatial concordance and greater specificity, yielding smaller, more interpretable marker sets. The same vector framework also extends to cluster- and gene-level co-location analysis.

**Availability and implementation:** jazzPanda is implemented as an open-source R/Bioconductor package, freely available at https://bioconductor.org/packages/jazzPanda. Analysis code for this article is at https://github.com/phipsonlab/jazzPanda_paper, with an accompanying analysis website at https://phipsonlab.github.io/jazzPanda_workflowr/. Datasets and scripts are deposited at https://zenodo.org/records/18149456.

## Introduction

Spatially resolved transcriptomics measures gene expression while retaining the physical location of cells and transcripts within a tissue, allowing the transcriptome to be studied in the context of tissue architecture. Early in situ approaches showed that RNA could be profiled directly in preserved tissue (Ke et al. 2013, Chen et al. 2015), and the introduction of spatial barcoding (Ståhl et al. 2016) was followed by rapid growth in both imaging-based and sequencing-based platforms. Commercial imaging-based technologies such as Xenium (Janesick et al. 2023), CosMx (He et al. 2022) and MERSCOPE (Yao et al. 2023) use fluorescence microscopy to localise transcripts from a predefined gene panel at subcellular resolution. Sequencing-based technologies such as Slide-seq (Rodriques et al. 2019), Visium and Visium HD (Oliveira et al. 2025), and Stereo-seq (Chen et al. 2022) capture broader transcriptome coverage, typically at a coarser effective resolution once the data are binned for analysis. A defining feature of imaging-based data is its sparsity: the transcript count per gene per cell is often zero, one or two (Figure 1a).

**Figure 1.**
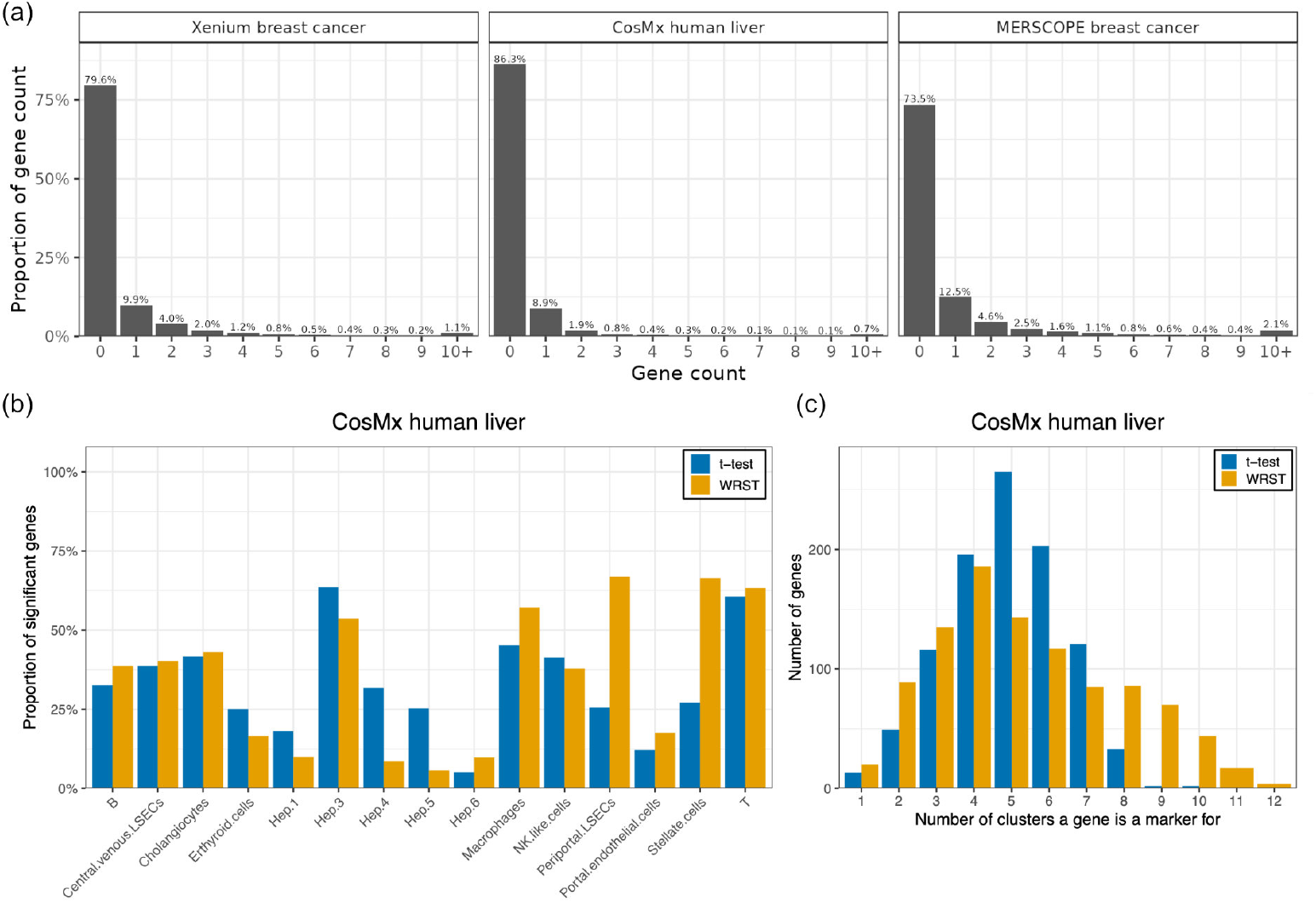
Sparsity of imaging based spatial transcriptomics data (a) and performance of single cell based marker analysis methods (b-c). (a) Barplot showing frequency of per cell gene counts in three different imaging based spatial transcriptomics datasets. (b) Proportion of significant marker genes for each cell type using either a t-test or Wilcoxon Rank Sum Test for a CosMx human liver dataset with a panel of 1000 genes. (c) Barplot showing how many clusters each gene is called a marker for, using either the t-test or Wilcoxon Rank Sum Test. It is relatively rare for a marker gene to be unique to only one cluster/cell type.

Cell type annotation is a central step in analysing this data, and most workflows still use tools developed for single-cell RNA sequencing (scRNA-seq). Cell types are commonly identified by clustering followed by marker gene detection, where markers are genes expressed more highly in a cluster than in the remaining cells. Common choices of test are the Wilcoxon rank sum test, the default in Seurat’s FindMarkers (Hao et al. 2021), and the t-test, available in Scanpy (Wolf et al. 2018). Both were designed for scRNA-seq and do not take account of the spatial coordinates of cells or transcripts. When applied directly to imaging-based data, these tests call a large proportion of the panel as significant for each cluster, and many of the genes returned are not specific to any one cell type (Figure 1b).

Spatial information is not absent from this part of the workflow. Several methods use the spatial coordinates directly for cell type clustering in spatial transcriptomics data, including CCST (Li et al. 2022), STELLAR (Brbić et al. 2022) and BASS (Li and Zhou 2022), which draw on the positions of neighbouring cells when assigning cell types or spatial domains. jazzPanda addresses a different and later step. We assume that cells have already been grouped into clusters, whether by a spatially aware method, such as BANKSY (Singhal et al. 2024), or a standard single cell method, for example Leiden clustering (Traag et al. 2019). We then ask which genes are spatially informative markers of those clusters. Clustering imaging-based data is itself a challenging task given the relatively small number of genes measured, and we do not attempt to solve it here; jazzPanda is designed to work with whatever clustering method a user prefers. Our aim is to incorporate the spatial locations of cells and transcripts into marker detection, the step immediately after clustering.

A related body of work detects spatially variable genes (SVGs), and it is worth being clear about how this differs from marker detection (reviewed in Yan et al. 2025). Global SVG methods such as SpatialDE (Svensson et al. 2018), SPARK (Sun et al. 2020) and SPARK-X (Zhu et al. 2021) test whether the expression of a gene is spatially structured across a tissue, without reference to cluster identity. A spatially structured gene need not mark any particular cluster, and a marker for a spatially dispersed cell type may show little global structure. A second group of methods, including C-SIDE (Cable et al. 2022) and CTSV (Yu and Luo 2022), tests whether a gene varies spatially within a given cell type. This is fundamentally different to the definition of a marker gene: a good marker is expressed consistently across the cells of its cell type; a gene that varies spatially within a cell type is not necessarily a marker gene for that cell type. SVG methods therefore ask whether a gene is spatially structured, whereas jazzPanda asks whether the spatial pattern of a gene is concordant with a particular cluster, a related but distinct question.

Here we present jazzPanda, a method for detecting spatially informative marker genes in imaging-based spatial transcriptomics. To overcome the sparsity of the data, jazzPanda divides the tissue into a grid of square, rectangular or hexagonal bins and sums the transcript detections and the cells of each cluster within each bin. This converts the two-dimensional coordinates of a gene, and of a cluster, into one-dimensional spatial vectors, and raises statistical power because the counts per bin are far larger than the counts per cell. Gene vectors can be built directly from transcript coordinates, so cell segmentation is optional; where segmentation is reliable, detections can instead be restricted to those assigned to cells. We define a marker as a gene whose expression is largely specific to a cluster and whose transcript detections overlap the locations of the cells in that cluster. jazzPanda measures this association in two complementary ways. For a single sample, permutation-based rank correlation tests whether a gene vector is associated with a cluster vector. For multiple samples, a lasso-regularised generalised linear model selects the clusters most relevant to each gene and also models sample-to-sample variability and the platform-specific background measured by negative-control probes. To our knowledge, jazzPanda is the only marker detection method to account jointly for spatial distribution, replication across samples, and platform background. We show by simulation that jazzPanda controls the false positive rate, apply both approaches to publicly available Xenium and CosMx datasets, and benchmark them against the Wilcoxon rank sum test and the t-test. The same framework extends to cluster- and gene-level co-location analysis which we demonstrate on a MERSCOPE dataset. jazzPanda is available as an open-source R/Bioconductor package (https://bioconductor.org/packages/jazzPanda).

## Results

### jazzPanda: a new method for spatially aware marker analysis

The foundation of our approach rests on binning transcript detections and cells into a grid across the tissue. Imaging-based spatial transcriptomics technologies can achieve sub-cellular resolution and provide the spatial coordinates of every detected transcript. However, the transcript counts per gene per cell tend to be very low, often zero, one or two counts (Figure 1a). Binning increases statistical power, because the counts per gene per bin are much larger than the counts at the individual cell level. Binning can also mitigate the effect of spatial autocorrelation at scales finer than the tile resolution. An ideal marker gene has higher and consistent expression in its target cluster than in the remaining clusters. In the spatial setting this means that a marker gene shows more transcript detections within the target cluster region than in other regions of the tissue. We therefore expect the binned transcript counts of a marker gene to increase with the number of cells of its target cluster across the tissue, so that the gene and cluster vectors are strongly associated. We assume that clustering has already been performed and that cluster labels are available for each cell, which allows jazzPanda to remain agnostic to the choice of clustering method. We propose two approaches to identify the genes whose vectors are strongly associated with a target cluster vector.

Figure 2 shows an overview of the jazzPanda method. We first discretise the tissue space into rectangular, square or hexagonal tiles, which converts the two-dimensional coordinates into one-dimensional spatial vectors. For each gene we define a gene vector (*g*_*i*_), and for each cluster a cluster vector (*x*_*i*_), both with length equal to the number of tiles.

**Figure 2.**
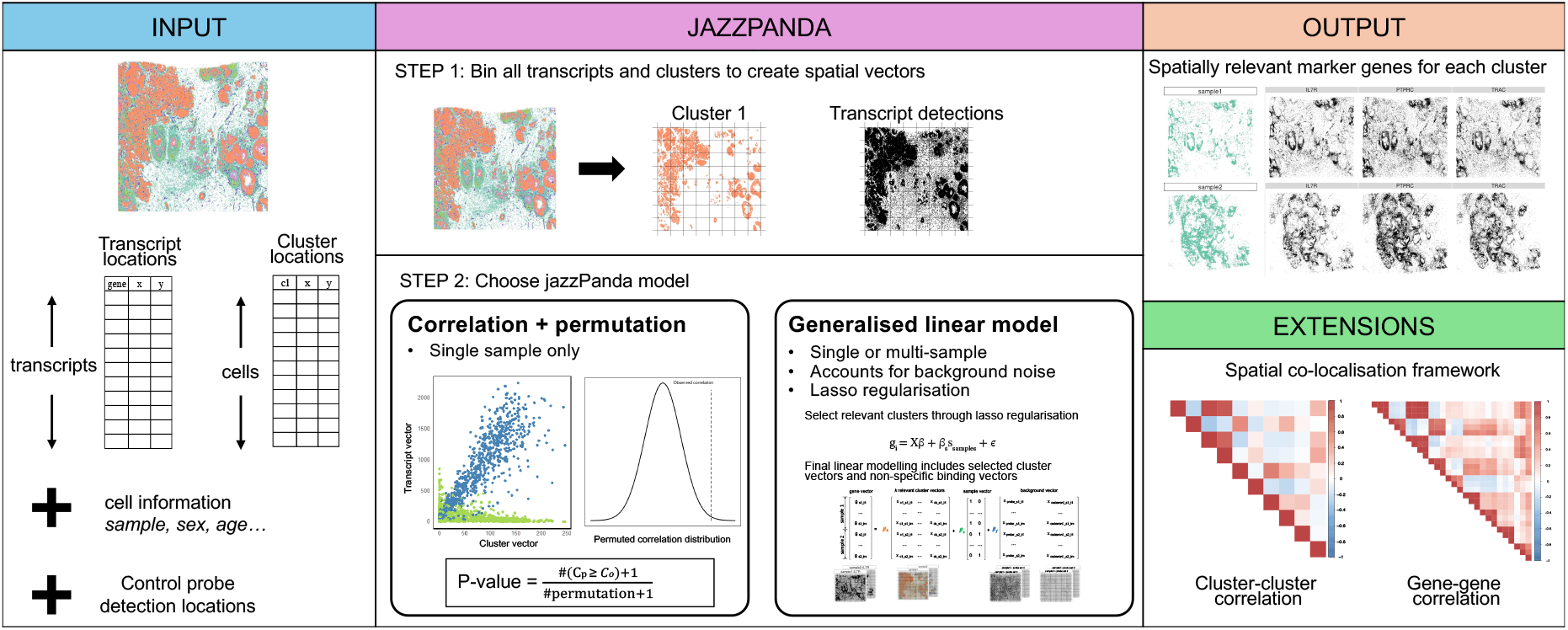
Overview of the jazzPanda approach. In Step 1, spatial vectors at the transcript and cluster level are created using square, rectangular or hexagonal bins. In Step 2, the spatial vectors are used as input for marker analysis using either a permutation framework to obtain statistical significance on the correlation estimates between clusters and gene vectors, or through a linear modelling approach.

The gene vector can be constructed in two ways. The first uses all transcript detections and is agnostic to how transcripts are assigned to cells. The second uses only transcripts that have been assigned to cells during segmentation, which excludes any detections that fall outside cell boundaries. jazzPanda provides both options, so that users can include or exclude out-of-cell transcripts depending on how reliable segmentation is for their data. In both cases the gene vector is obtained by summing the relevant transcript counts within each tile. The cluster vector is obtained by counting the number of cells from that cluster in each tile, and therefore always depends on the segmentation and clustering that have already been performed. We define the marker genes of a cluster as the genes whose gene vector is strongly associated with the target cluster vector.

Once the spatial vectors have been constructed, they can be used as input for standard statistical techniques. We provide two approaches for detecting marker genes. The first is based on correlation and the second on linear modelling. Both approaches are described below, with further details given in ‘Methods’.

### jazzPanda-correlation: permutation testing approach

We use the Spearman rank correlation to measure the strength of the monotonic association between a gene vector and a cluster vector. Spearman correlation is less sensitive to outliers than Pearson correlation, which is why we adopt it here. A gene whose vector has a strong positive correlation with a target cluster vector is assigned as a marker gene for that cluster. We obtain a significance level for each correlation using permutation. In each permutation, we randomly shuffle the cluster labels across the cells, rebuild the cluster vectors from the shuffled labels, and recalculate the correlation between every gene vector and every shuffled cluster vector. We repeat this process a number of times, for example 10,000 permutations. The p-value for an observed correlation is calculated as the proportion of permutations in which the correlation is at least as large as the observed value, using the equation in Phipson & Smyth (Phipson and Smyth 2010). The permutation p-values are then adjusted for multiple testing using the Benjamini and Hochberg method (Benjamini and Hochberg 1995).

### jazzPanda-glm: generalised linear modelling approach

A limitation of the correlation approach is that it cannot account for additional covariates in a straightforward way. When samples are jointly clustered so that clusters are consistently labelled, the model should account for multiple samples rather than analysing each sample separately. We treat the gene vector as the response variable and the cluster vectors as the predictor variables, and fit a linear model for each gene. Because the original count scale faithfully preserves the linear relationship between the cluster and gene vectors, we apply no further scaling or transformation.

Each gene is associated with a different subset of clusters, so we use lasso regularisation to select, for each gene **g**_*i*_, the clusters that are most relevant (Friedman et al. 2010, Tay et al. 2023). The lasso penalty parameter *λ*_*i*_ is estimated separately for each gene by cross-validation, because genes differ in their expression levels (see ‘Methods’ for further details). This yields a set of clusters with non-zero coefficients for each gene, which are then used to fit a final linear model. To account for study designs with multiple samples, we add a sample-level vector (**S**) to the final model. Including this term accounts for sample-to-sample variability and reduces sample-specific effects, so that the model identifies marker genes that are shared across samples.

The output of the final model can be interpreted in two ways. To identify unique marker genes, those that mark a single cluster, we assign each gene to the cluster with the smallest p-value and the largest positive coefficient. In this setting, clusters effectively compete for assignment by maximising how well they explain the gene expression vector. To identify shared marker genes, those that mark more than one cluster, we report all clusters that are significant for the gene. This second criterion is less stringent, with the lasso regularisation enforcing sparsity and retaining only the most informative clusters.

### Accounting for non-specific binding of RNA molecules

A further advantage of the linear modelling approach is that it can account for background noise, by including additional covariates that represent unwanted variation. Spatial platforms implement different negative controls to assess assay quality, and these fall into two broad types. The first type consists of barcode sequences that do not match any real probe in the assay. All three platforms include controls of this type: negative control codewords on Xenium, falsecodes on CosMx, and blank genes on MERSCOPE (Figure 3a-c, Supplementary Figure S1). The second type consists of negative probes, which are artificial sequences that are absent from the study organism and are used to quantify non-specific binding. Xenium and CosMx also include negative probes (10x Genomics 2024, Beechem et al. 2023).

**Figure 3.**
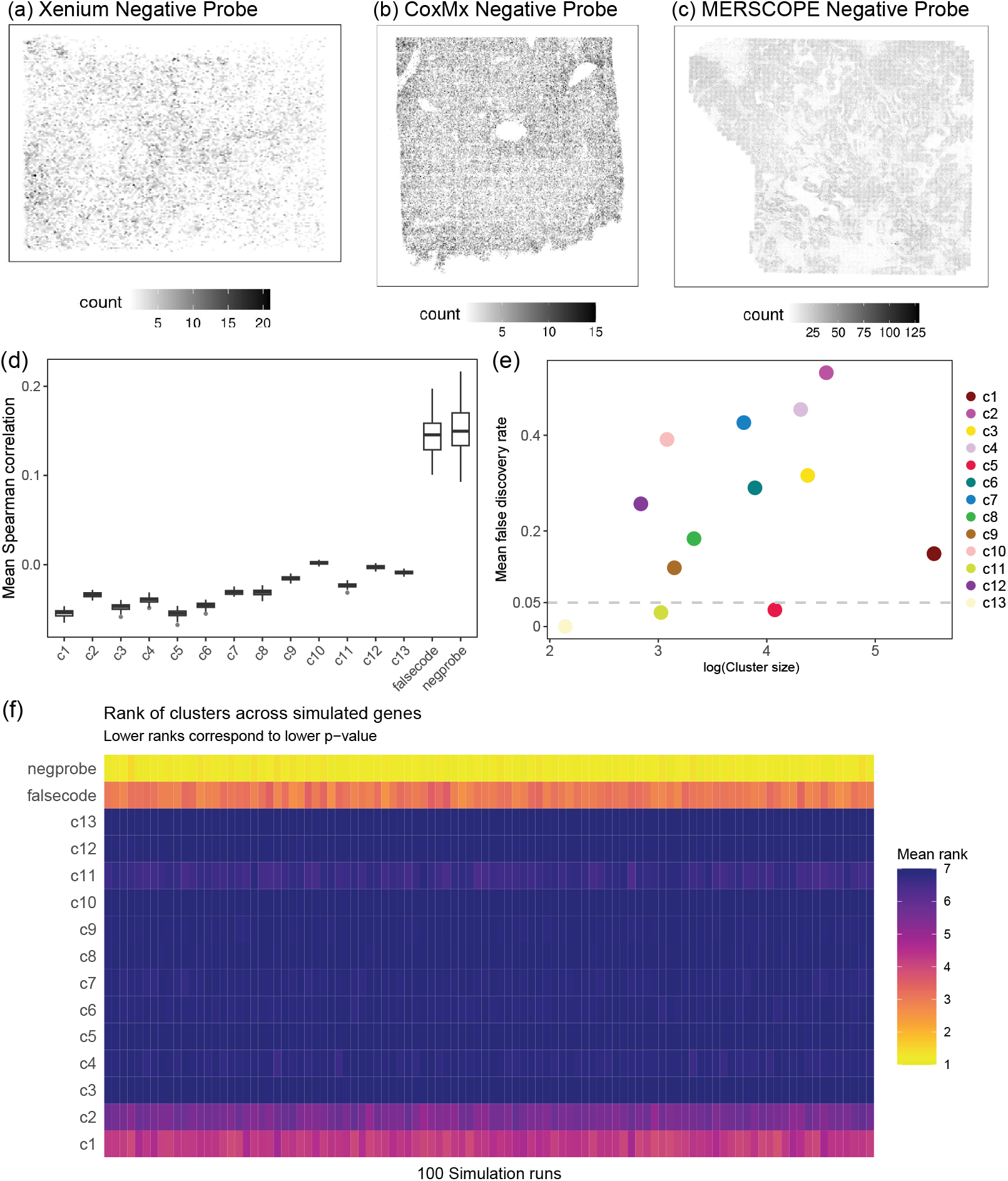
Spatial distribution of negative control probes on imaging based platforms (a)-(c) and evaluating jazzPanda’s performance using simulated datasets (d)-(f). (a) Negative control probes for the Xenium human breast cancer sample. (b) Negative control probes for the CosMx healthy human liver sample. (c) Negative control probes for the MERSCOPE human breast cancer sample. (d) Boxplot of the Spearman correlation between uniformly generated gene vectors, negative control vectors, and cluster vectors, across 100 datasets each containing 100 uniformly generated genes. (e) Scatter plot of the relationship between cluster size and the mean false positive rate from jazzPanda-correlation, calculated without a correlation cut-off. For each dataset the false positive rate is the proportion of the 100 null genes called significant (p-value *<* 0.05), averaged across the 100 simulated datasets. (f) Mean ranking of cluster and negative control vectors derived from the gene-wise p-values using the linear model approach across each simulation run.

jazzPanda represents this background using the same binning strategy. For each category of negative control, we sum the corresponding detections within each tile to form a background noise vector (f). For Xenium, for example, this produces two background vectors, one for the negative control codewords and one for the negative probes. These vectors are added as covariates to the final linear model, so that the model accounts for platform-specific background and improves control of the false discovery rate.

### Type I error rate control: null simulations

To evaluate the two approaches when no true marker signal is present, we simulated uniform gene and negative control vectors based on the CosMx human liver cancer dataset, which carries the largest gene panel in our study (1,000 genes) and therefore offers the widest pool of genes to sample from (Supplementary Figure S2). We randomly selected 100 real genes, and for each gene we generated the same number of uniformly distributed points as its observed transcript detections, placed at random across the tissue space. For each negative control category we generated uniformly distributed points equal in number to its observed detections. We then applied jazzPanda to each simulated dataset, using the real cluster vectors from the liver cancer dataset, to test whether the uniform genes would be assigned as marker genes to any cluster.

Figure 3d shows the distribution of the Spearman correlations of the simulated gene vectors with the real cluster vectors and with the negative controls across the 100 datasets. The uniform gene vectors have almost no correlation with the real clusters, as expected (median correlation −0.028). Their correlation with the simulated falsecode and negprobe background vectors is higher, at 0.16 and 0.198 respectively. Because the gene-cluster correlations are so low, a correlation cut-off on its own calls none of the uniform genes as markers. The permutation p-values behave differently: with 10,000 permutations, the proportion of significant genes ranged from close to 0% to more than 40% across the clusters, with the average Type I error rate only controlled at 5% for clusters 5, 11 and 13 (Figure 3e). In practice it is therefore important to examine the correlation alongside the permutation p-value rather than relying on the permutation p-value alone. A Spearman correlation cut-off as small as 0.05 removes these false positives, reducing the false positive rate to below 1% for all clusters (Supplementary Figure S3).

The permutation approach cannot easily incorporate background signal when calculating p-values, whereas the linear modelling approach can include the background vectors directly. When a gene’s spatial pattern is highly correlated with the background, these vectors absorb the signal, so the linear model does not assign the gene to any cluster. Figure 3f shows the ranking of the clusters and negative controls based on the final linear model coefficients for 100 simulated datasets. The negprobe or falsecode vector is ranked as the most relevant term in 89.27% and 10.73% of cases respectively, so the top term by minimum p-value and largest coefficient is always a negative control and the gene is not assigned to any cluster. The lasso step can still retain additional real clusters, most often one (35.68%) or two (40.99%) and only rarely three (3.95%) or four (1.44%). These tend to be large clusters or clusters with more uniformly distributed cells. Even when extra clusters are retained, the background remains the top-ranked term after ordering by minimum p-value and largest coefficient.

Taken together, these simulations show that both approaches can control false positives, and that they are complementary in practice. jazzPanda-correlation is reliable when the permutation p-value is examined together with a correlation cut-off, and a correlation threshold as small as 0.05 keeps the false positive rate below 1%. jazzPanda-glm additionally models platform-specific background, which makes it well suited to data where non-specific binding is a concern, as well as when multiple samples need to be analysed together.

### jazzPanda identifies specific, spatially coherent marker genes in real spatial datasets

We illustrate jazzPanda on two real publicly available datasets that differ in panel size and experimental design: a single CosMx healthy human liver sample (1,000 genes, 332,877 cells, 15 annotated cell types), and a Xenium HER2+ human breast cancer dataset with two biological replicates (313 genes, nine shared clusters). We use the liver sample as a single-sample example, and the breast cancer dataset to show how jazzPanda-glm accounts for multiple samples and platform-specific background. In both cases we used 40 × 40 square bins to construct cluster vectors from the cell coordinates and gene vectors from the transcript detection coordinates.

We first applied jazzPanda-correlation to the liver sample, where the annotated cell types range from abundant hepatocytes to rarer immune and endothelial populations (Figure 4a). Supplementary Figure S4 shows the top three markers for every cluster. Taking stellate cells as an example (Figure 4b), jazzPanda-correlation identified 130 marker genes with a significant adjusted permutation p-value and an observed Spearman correlation above the per-cluster cut-off. The top marker was insulin-like growth factor-binding protein 7 (IGFBP7), with a Spearman correlation of 0.904 between the gene and cluster vectors and an adjusted permutation p-value of 0.0018 (Figure 4c). IGFBP7 is differentially expressed in hepatic stellate cells and is used as a specific marker for this population in single cell human liver studies (Payen et al. 2021). The IGFBP7 transcripts are spatially concordant with the stellate cell cluster across the tissue (Figure 4b, c), and the gene and cluster vectors show a clear monotonic relationship (Supplementary Figure S5).

**Figure 4.**
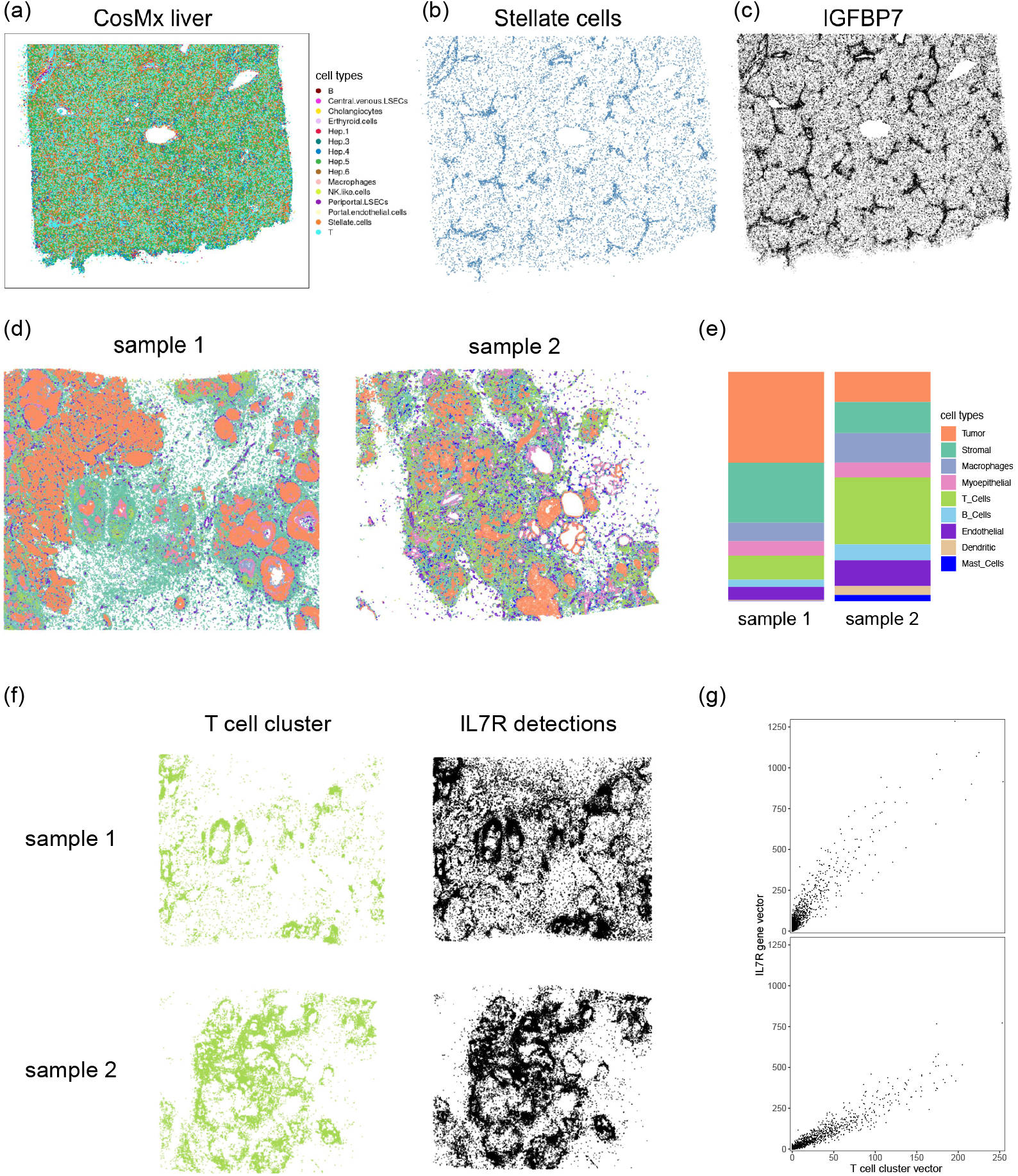
Application of jazzPanda to CosMx healthy liver sample (a-c) and Xenium HER2+ human breast cancer samples (d-g). (a) Spatial visualization of cell type coordinates (b) Spatial visualization of cell type coordinates for the Stellate cell cluster (c) Spatial visualization for the top marker gene IGFBP7 using jazzPanda-correlation (d) Spatial visualization of cell type coordinates for each sample, where each point represents a cell and is coloured by cell type (e) Cell type composition per sample (f) Spatial visualization of cell type coordinates for the T cell cluster and the corresponding transcript detections for the top marker gene, IL7R, selected by the linear modelling approach (d) The relationship between the T cell cluster vector (x axis) and the marker gene vector (y axis), each point refers to one grid value. Top panel is for sample 1 and bottom panel is for sample 2.

Applying jazzPanda-glm to the same sample with background detections included in the linear model, also returned IGFBP7 as the most significant marker for stellate cells. jazzPanda-glm identified 40 marker genes for stellate cells, far fewer compared to the correlation approach. Across clusters, the top three markers were broadly concordant between the two approaches, with at least one gene in common for most clusters, although the ranks and some individual genes differed (Supplementary Figures S4, S6). Again, the cluster and gene vectors showed a clear monotonic relationship for jazzPanda-glm marker genes (Supplementary Figure S7).

To demonstrate jazzPanda-glm on a multi-sample design with background detections accounted for, we used the Xenium breast cancer dataset, which contains two biological replicates with nine shared clusters. The two samples have distinct spatial organisation and cell type composition (Figure 4d, e). We converted the two types of negative probes into two background vectors with the same binning dimensions and included them, together with a covariate for sample, in the final linear model (Figure 3a, Supplementary Figure S1a, b). Using T cells as an example, jazzPandaglm identified 31 marker genes, and interleukin 7 receptor (IL7R) as the top marker. The T cell clusters occupy distinct spatial patterns in the two samples, yet within each sample the spatial concordance between the IL7R transcripts and the T cell cluster is high, and the gene and cluster vectors show a clear linear relationship in both replicates (Figure 4f, g). IL7R and its ligand IL7 are required for T cell development (Wang et al. 2022), and higher IL7R expression has been associated with more aggressive breast tumours (Al-Rawi et al. 2004). The top three markers for all nine clusters are shown in Supplementary Figure S8 and the linear relationship between the cluster and gene vectors are shown in Supplementary Figure S9.

Across both datasets, jazzPanda returns markers that are spatially concordant with their clusters and recover known cell type markers. On the single liver sample the correlation and linear modelling approaches agree on the top markers, and on the Xenium data the linear model identifies shared markers across replicate samples while accounting for platform background within a single model.

### Comparison of jazzPanda with single cell marker analysis methods

A common approach for marker gene analysis in single cell data is the Wilcoxon Rank Sum Test comparing each cluster to the remaining clusters, implemented in the ‘FindAllMarkers’ function in Seurat, as well as an implementation in Scanpy. A recent benchmarking study found that simple tests such as the Wilcoxon Rank Sum Test and the t-test often outperform more complex methods in single cell data (Pullin and McCarthy 2024). A drawback of the Wilcoxon Rank Sum Test is that it is non-parametric and cannot account for additional covariates such as sample or sex. We compared the marker genes detected by jazzPanda, the Wilcoxon Rank Sum Test, and the moderated t-test implemented in the limma R package. The ‘FindAllMarkers’ function applies a default log fold-change cut-off of 0.1. We removed this cut-off so that the statistical tests are compared without additional filtering. For the moderated t-test and jazzPanda-glm we included a sample-level covariate for multi-sample datasets.

We compared the three methods on the CosMx healthy liver and Xenium breast cancer datasets (Figure 5). For each dataset we selected a large, a medium and a small cluster to check for any bias related to the number of cells in a cluster. We defined marker genes as those genes that had a positive log-fold change and a Benjamini and Hochberg adjusted p-value below 0.05 for both the Wilcoxon Rank Sum Test and the moderated t-test. For jazzPanda-correlation and jazzPanda-glm we selected marker genes as described above.

**Figure 5.**
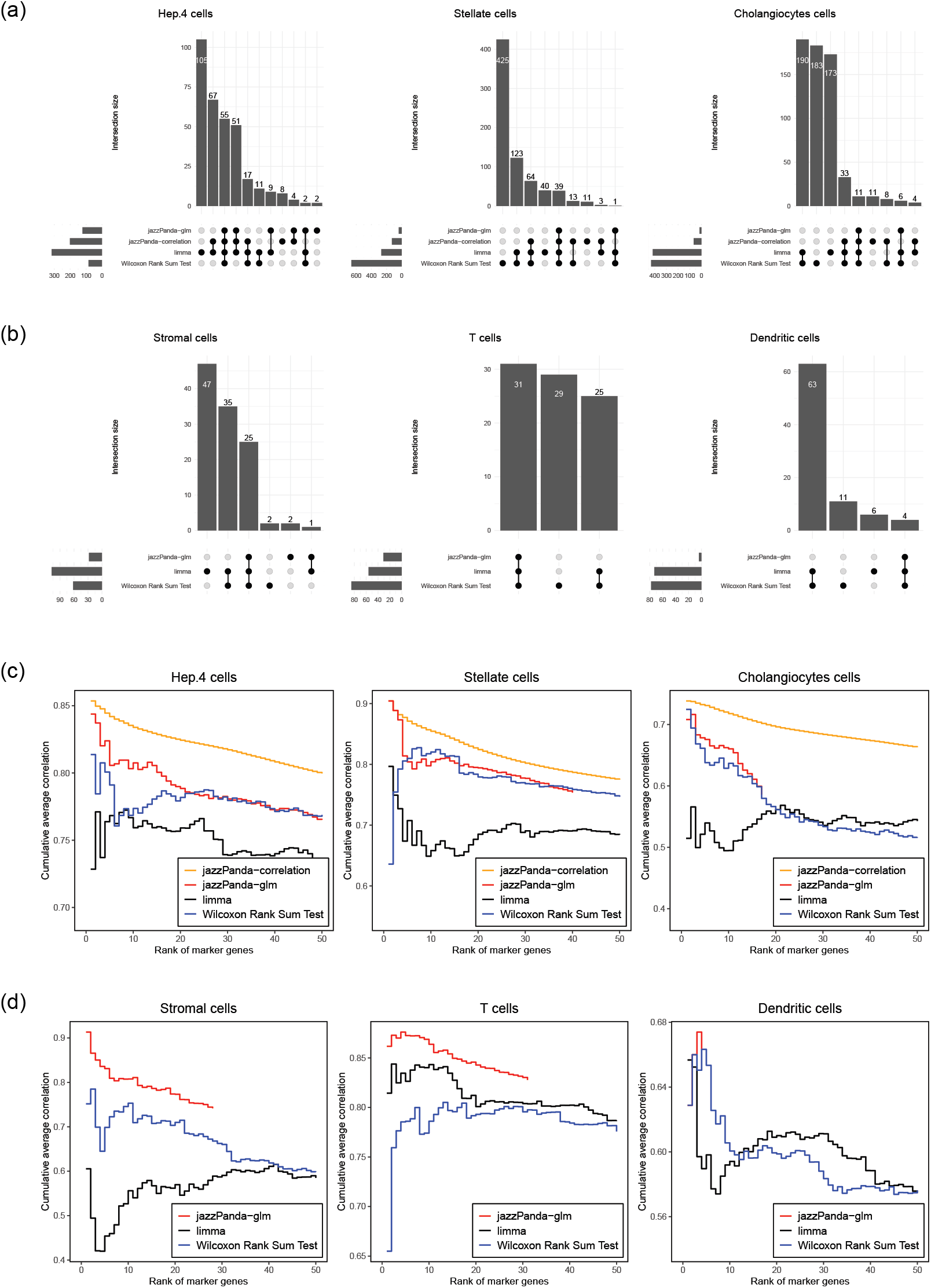
Comparison of marker analysis methods. (a) Upset plots showing the overlap of genes detected by each method for a large cluster (Hep.4 cells), a medium cluster (stellate cells), and a small cluster (cholangiocytes) in the CosMx healthy liver sample. (b) Upset plots showing the overlap of genes detected by each method for a large cluster (stromal cells), a medium cluster (T cells), and a small cluster (dendritic cells) in the two Xenium breast cancer samples. (c) Cumulative average correlation of the top 50 ranked marker genes for each method in the healthy liver sample. (d) Cumulative average correlation of the top 50 ranked marker genes for each method in the two breast cancer samples.

Both jazzPanda approaches identified far fewer marker genes per cluster than the Wilcoxon Rank Sum Test and the moderated t-test, and the jazzPanda marker sets were largely subsets of them (Figure 5a, b). The number of marker genes called by jazzPanda tended to scale with cluster size, with more markers for larger clusters and fewer for smaller ones, an effect that was more pronounced for jazzPanda-glm. On the liver sample jazzPanda-correlation also called some genes that no other method returned (Figure 5a); some of these may reflect non-specific binding, the signal that jazzPanda-glm removes to return a smaller and more specific set. Upset plots for all clusters are shown in Supplementary Figures S10 and S11.

Although the two standard tests return large marker sets, their top-ranked genes may still be biologically relevant. We therefore assessed how spatially coherent each method’s rankings were, by calculating the spatial correlation between the ranked marker genes and the target cluster and plotting the cumulative mean correlation of the top 50 markers for each method (Figure 5c, d). This metric favours jazzPanda-correlation by construction, since it ranks genes on exactly this quantity, and it has the highest cumulative correlation on the single sample (Figure 5c). More informatively, jazzPanda-glm, which ranks by model coefficient rather than correlation, also concentrated high-correlation genes near the top of its list, with its top gene usually strongly correlated with the cluster (Figure 5c, d). The Wilcoxon Rank Sum Test and the moderated t-test were more variable, with top-ranked genes that were highly correlated for some clusters but not others. Supplementary Figures S12 and S13 show the cumulative correlation plots for all clusters. Taken together, jazzPanda returns smaller, spatially coherent marker sets that are subsets of those from the standard tests, and jazzPanda-glm does so while accounting for background and multiple samples.

### Extension of statistical framework: spatial co-location

The binning step that underpins jazzPanda produces spatial vectors for both transcripts and cell types, and these vectors can be reused for analyses beyond marker detection. Correlating the cluster vectors with one another measures how strongly cell types co-locate across a tissue, giving an estimate of its global spatial organisation. A positive correlation between two clusters indicates that the cell types tend to occupy the same regions, while a negative correlation indicates that they occupy distinct regions.

We demonstrate this on a MERSCOPE human breast cancer sample (Figure 6a) by computing the pairwise Spearman correlation between all cluster vectors (Figure 6b). The strongest co-location is among the tumour clusters, with Tumor_luminal_ERBB2 correlating with Tumor_IFNg_APC and Tumor_Cycling_G2M at 0.96 and 0.9 respectively. The immune compartment shows its own structure, with TAM_C1QC and activated T cells co-locating (Spearman correlation 0.66). Negative correlations are modest across the sample, the largest being between TAM_SPP1 and endo_blood at −0.22. These relationships are borne out in the spatial cluster patterns (Supplementary Figure S14), where the positively correlated tumour clusters show clearly overlapping distributions, and the negatively correlated clusters occupying distinct regions of the tissue space.

**Figure 6.**
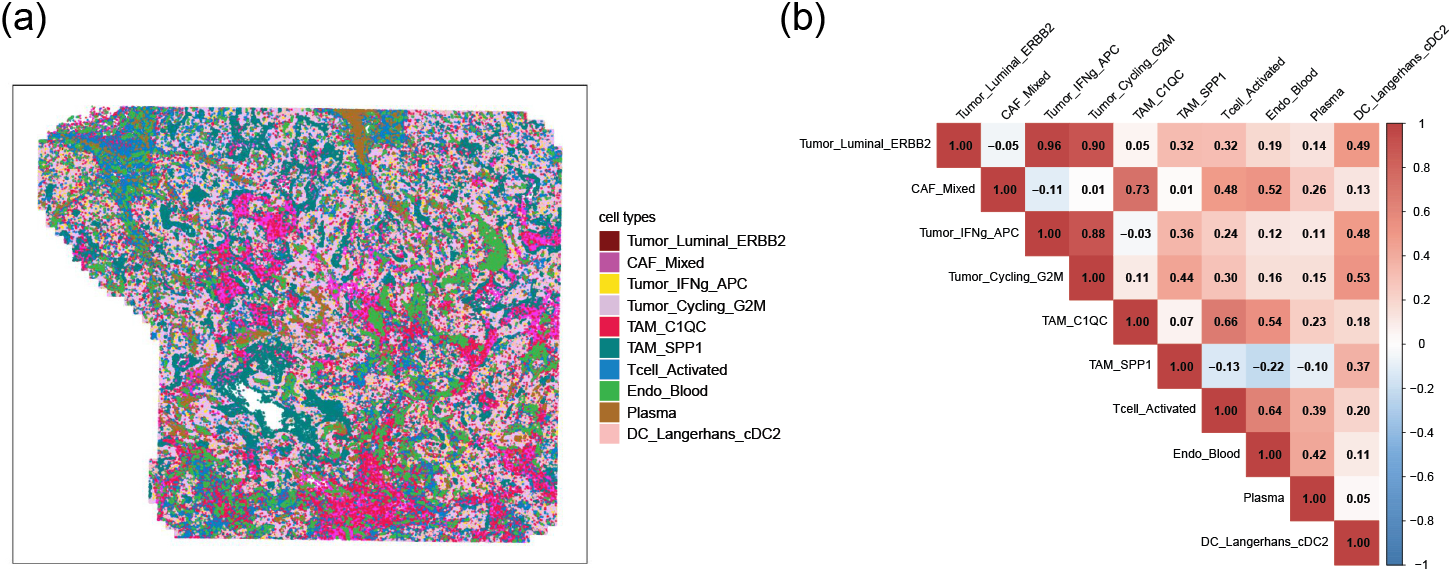
Extension of the jazzPanda framework. (a) Spatial visualization of cell type coordinates for a MERSCOPE human breast cancer sample, where each point represents a cell and is coloured by the annotated cell type. (b) Pairwise Spearman correlation between the cluster spatial vectors.

The same framework applies to gene-gene correlation, where a positive correlation identifies genes co-expressed at the same locations and a negative correlation identifies genes that are spatially exclusive. As a proof of concept, the pairwise Spearman correlations among the top three markers of each cluster recapitulate the cluster co-location structure (Supplementary Figure S15, Supplementary Table T1). Because every gene has a spatial vector, the same approach could be applied across the full panel to identify spatially co-expressed gene modules, for example through a spatial adaptation of weighted correlation network analysis (Zhang and Horvath 2005).

## Technical performance

### Effect of grid size on marker gene detection

The grid size trades spatial resolution against statistical power and is a parameter the user must set, so we assessed how robust jazzPanda-glm marker detection is to this choice. We compared the markers detected across square grids from 10 × 10 to 100 × 100 for a large, a medium and a rare cluster in the Xenium human breast cancer dataset (Figure 7a). In general jazzPanda-glm is robust to grid size, with the largest intersection of genes shared by every grid size in the upset plot. For this particular dataset, 10×10 may be too coarse, as it missed marker genes detected at the other resolutions. The result for every cluster is shown in Supplementary Figure S16.

**Figure 7.**
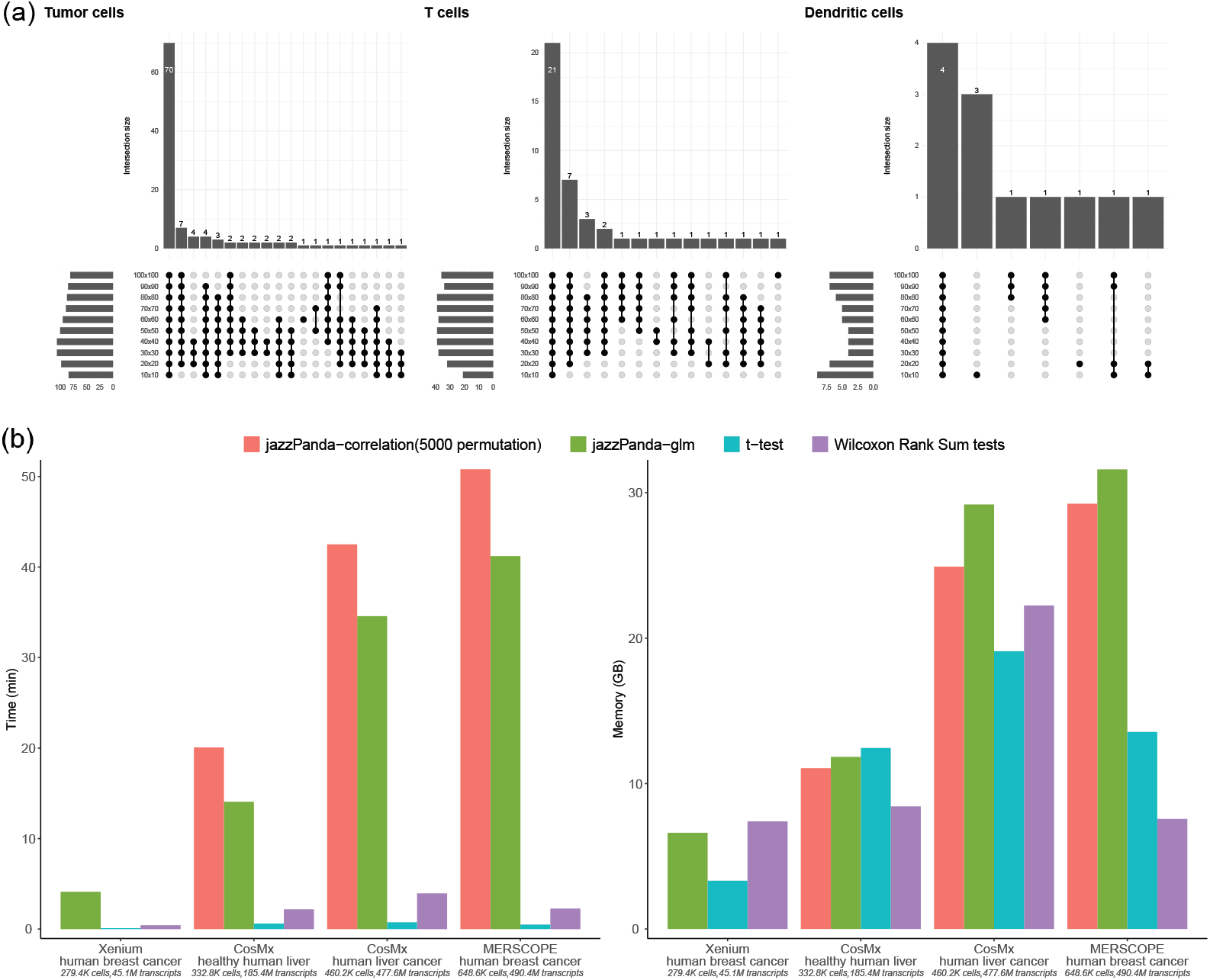
Technical performance of jazzPanda. (a) Upset plot showing the consistency of the marker genes identified by jazzPanda-glm across grid sizes from 10^2^ (10×10) to 10^4^ (100×100), for three clusters of varying size in the Xenium human breast cancer dataset: a large cluster (Tumour cells), a medium cluster (T cells) and a rare cluster (dendritic cells). (b) Runtime and peak memory usage for marker gene detection across four datasets (Xenium human breast cancer, CosMx healthy human liver, CosMx human liver cancer and MERSCOPE human breast cancer), comparing jazzPanda-correlation (5000 permutations), jazzPanda-glm, limma and the Wilcoxon Rank Sum Test. The two jazzPanda methods were parallelised across five cores.

We also compared the top markers for each cluster across the range of grid sizes using the mean pairwise Jaccard index. Agreement was high for clusters with well-populated marker sets (0.73 to 0.85), indicating the recovered markers are largely stable to the choice of grid size. Lower values occurred only for clusters with very few markers, where the index is sensitive to individual genes (Supplementary Table T2).

As a practical guide for selecting the grid size, we recommend a grid size such that the average number of cells per tile is at least one across all clusters. For example, if the smallest cluster contains 100 cells, the length of the grid should be at most 100, which could be represented by 10×10 square grids. This also keeps the spatial vectors from becoming too sparse, so that a reliable lasso penalty can be estimated for each gene.

### Computational complexity

We measured the time and peak memory needed to detect marker genes with each method across four datasets, which span a range of panel sizes and cell and transcript counts (Figure 7b), measured on an HPC system with 12 cores and 300 to 400 GB of memory. jazzPanda-correlation is the slowest, at roughly 20 to 50 minutes, owing to the permutation step, and jazzPanda-glm takes around 40 minutes for the largest dataset (MERSCOPE human breast cancer). The Wilcoxon Rank Sum Test and moderated t-test are much faster, at 10 seconds to 4 minutes.

For jazzPanda, constructing the spatial vectors is the most time-consuming step, and its cost is driven mainly by the number of transcript detections (Supplementary Figure S17). Because each vector is built independently, this step runs in parallel, so additional cores reduce the total time. Marker detection with jazzPanda-glm takes under five minutes for the smallest dataset, and on five cores both jazzPanda approaches completed the MERSCOPE human breast cancer sample, which contains over 490 million transcripts, within 50 minutes. Memory use rises with dataset size and is highest for jazzPanda-glm, at roughly 8 to 30 GB (Figure 7b). Hexagonal bins represent spatial information slightly more faithfully than square bins but are more expensive to generate (Supplementary Figure S18). Square bins offer a good trade-off and we used them throughout our analysis.

## Discussion

Cell type annotation is a key step in the analysis of spatial transcriptomics data. We have developed two complementary approaches for detecting spatially informative marker genes and implemented them in the jazzPanda Bioconductor R package, and we have applied them to publicly available CosMx, Xenium and MERSCOPE datasets. Because gene vectors can be built directly from transcript coordinates, jazzPanda does not require transcripts to be assigned to cells, which draws on more of the available signal than cell-level expression alone. Where transcript coordinates are not available, the gene vectors can instead be built from the cell-level count matrix.

The simulations show that accounting for the background signal measured by negative-control probes reduces the false positive rate. This is a particular strength of jazzPanda-glm, which can model the background together with sample-to-sample variation, and for this reason we recommend it for designs with multiple samples; the correlation approach offers a simpler alternative for a single sample. Compared with the Wilcoxon rank sum test and the moderated t-test, jazzPanda returns a smaller and more spatially specific set of marker genes for each cluster. A shorter and more distinctive marker list is intended to make manual cell type annotation more straightforward.

jazzPanda works best when clusters are spatially separable, particularly when the aim is to find markers unique to a cluster rather than markers it shares with others. If the data are over-clustered, merging similar clusters can improve the results, and the cluster-cluster correlations returned by jazzPanda can indicate when clusters overlap in space. The size of a cluster can also affect power, particularly for jazzPanda-glm: across all datasets larger clusters tended to yield more significant markers than smaller ones. The correlation approach is limited in that it cannot adequately account for background signal or additional covariates, such as sample or sex.

Two practical choices affect the analysis. Hexagonal bins can capture spatial structure slightly more faithfully than square or rectangular bins, but at a significantly higher computational cost, and in our experience square or rectangular bins performed well. For grid size, a useful rule of thumb is to choose bins such that the average number of cells per bin is at least one in every cluster; this also stabilises the gene-wise penalty estimates and helps to satisfy the linearity assumption of the model. Building the gene and cluster vectors is the most time-consuming step of the pipeline, but once they have been computed the modelling step is fast.

Because jazzPanda depends only on a grouping of cells, it is compatible with any clustering method and can equally be applied to niches or user-defined regions of interest rather than clusters alone. The same vector framework supports further analysis, including the co-location of clusters and genes, and could be extended to spatially informed ligand-receptor analysis and weighted gene co-expression networks. In summary, jazzPanda provides a spatially aware approach to marker analysis in imaging-based spatial transcriptomics that produces specific, spatially coherent markers while controlling the false positive rate, and it is freely available as a Bioconductor R package.

## Methods

### Statistical model framework for jazzPanda

For a given spatial dataset we profile *n* genes in each sample *s* ∈ {1, …, *S*}. Each gene *i* has a number of *d*_*is*_ transcript detections profiled across the tissue sample *s*. Each *d*_*is*_ has an *x* and *y* coordinate. We divide the spatial domain into a grid of bins *V* ^1^, …, *V* ^*h*^, …, *V* ^*v*^, which may be square, rectangular or hexagonal, and use the same grid for all genes and clusters.

For each gene *i* ∈ {1, …, *n*} in sample *s* we count the detections falling in each bin to form a gene vector **g**_*is*_ ∈ ℝ^*v*^, where each element 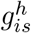 of the gene vector represents the sum of the detections of gene *i* within the boundary of bin *V* ^*h*^ in sample *s*

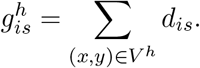

Using the same grid, we count the number of cells of cluster *j* ∈ {1, …, *c*} in sample *s* in each bin to form a cluster vector **x**_*js*_ ∈ ℝ^*v*^, where

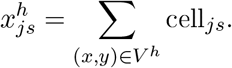

Stacking across samples, we define the gene matrix **G** and design matrix **X**:

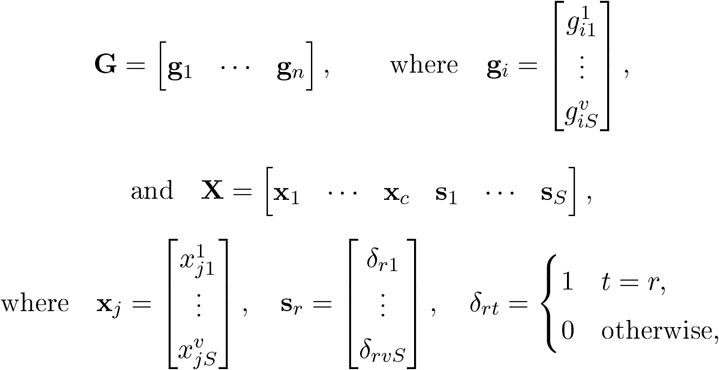

where **s**_*r*_ is an indicator vector for sample *r*.

Each gene vector **g**_*i*_ is modelled as

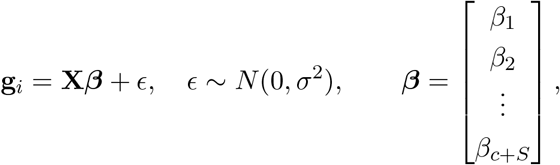

where the coefficients ***β*** are estimated by lasso regression with the penalty *λ*_*i*_ chosen by crossvalidation:

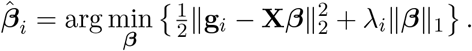

The *k* clusters with non-zero coefficients are retained. For each gene we then fit a final model on the selected cluster vectors together with the sample vectors **s** and the background vectors **f** :

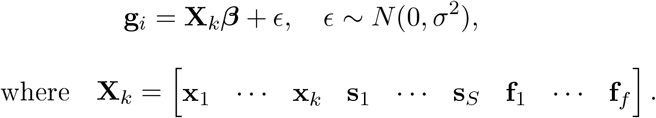

For a unique marker, gene *i* is assigned to the cluster with the smallest p-value and the largest coefficient.

### Correlation approach for a single sample

For a single sample, jazzPanda tests the association between each gene vector and each cluster vector using Spearman rank correlation. For gene *i* and cluster *j* we compute the observed Spearman correlation *ρ*_*ij*_ between **g**_*i*_ and **x**_*j*_. A permutation null is generated by randomly shuffling the cluster labels across cells and rebuilding the cluster vectors from the permuted labels, while holding the gene vectors fixed. Recomputing the correlations under *B* permutations yields a null distribution for each gene-cluster pair, from which a permutation p-value is obtained and adjusted across genes using the Benjamini-Hochberg procedure. Gene *i* is called a marker of cluster *j* when its adjusted p-value is below 0.05 and its observed correlation exceeds a cut-off. We used a cut-off of 0.05 for the simulation; for the real datasets the cut-off was set per cluster to the 75th percentile of the observed correlations.

### Estimation of the lasso penalty parameter

Given the bin values for a gene vector and the cluster vectors, we use stratified sampling to divide the bins into ten homogeneous folds that represent the overall distribution (Lohr 2021). A Gaussian linear model is trained on nine folds and validated on the tenth, repeated so that each fold serves once as the validation set. We select the conservative *λ*_1*se*_, the largest penalty within one standard error of the minimum cross-validation error, to avoid over-fitting.

### Datasets and implementation

This study used four publicly available datasets from three platforms (Xenium, CosMx and MERSCOPE); full URLs and accession details are given in the Data Availability section. All analyses were performed in R (4.5.1), and the package versions are recorded in the “Session Information” of each script at https://phipsonlab.github.io/jazzPanda_workflowr/. Dataset-specific quality control and clustering are described in Supplementary Methods.

### Simulation

To evaluate the false positive rate (Type I error), we used the CosMx human liver cancer data, the largest single sample dataset. We selected 100 genes from the panel at random and, for each, simulated a null spatial pattern: transcripts were placed on a 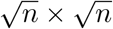 grid spanning the tissue window, where *n* is the number of detections for that gene, with independent uniform jitter added to each coordinate to avoid the local clustering of fully random placement. Background control detections were generated in the same way. We then applied jazzPanda-correlation and jazzPanda-glm on a 40 × 40 grid, repeating the procedure 100 times.

### Running jazzPanda

Spatial vectors were created with *get_ vectors* using square bins: 40 × 40 for Xenium human breast cancer, CosMx human healthy liver and CosMx human liver cancer, and 50 × 50 for MER-SCOPE human breast cancer. The linear modelling approach (*lasso_ markers*) was applied to every dataset, and the correlation approach (*compute_ permp*) to the single-sample datasets, using 5,000 or 10,000 permutations. Background vectors were built from the negative controls with *create_ genesets* and binned on the same grid as the genes and clusters: negative-control probe and codeword vectors for Xenium, negative-probe and falsecode vectors for CosMx, and a blank-gene vector for MERSCOPE.

### Wilcoxon rank sum test

Marker genes were identified with the *FindMarkers* function in Seurat (5.3.0) and selected at an adjusted p-value below 0.05.

### limma

We used moderated t-tests from the limma package to demonstrate marker analysis with t-tests. Counts were normalised with *normCounts* from the speckle (1.8.0) R package. A design matrix was specified with one group per cluster, and a contrast matrix was built to compare each cluster with the average of all others. The limma (3.64.1) pipeline was applied (*lmFit, contrasts. fit and eBayes*) and the results summarised with *decideTests*.

### Robustness and computational performance

The effect of bin length on marker selection was assessed for the two-sample Xenium human breast cancer data across grids from 10 × 10 to 100 × 100; the full procedure is given in Supplementary Methods. To quantify the stability of marker detection across grid sizes, we identified the set of top marker genes for each cluster at each grid size from 10 × 10 to 100 × 100, computed the Jaccard index (the size of the intersection divided by the size of the union) between the marker sets for every pair of grid sizes, and report the mean over all pairs for each cluster. Runtime and peak memory were profiled on the WEHI HPC cluster with the *peakRAM* R package (1.0.2); the computational complexity experiments, with respect to the number of detections and to bin shape and length, are described in Supplementary Methods.

## Data availability

All raw and processed datasets used in this study are publicly available via Zenodo https://zenodo.org/records/18149456. The raw data can be accessed directly from the following sources:

- 10x Xenium Human HER2+ breast cancer data: https://www.10xgenomics.com/products/xenium-in-situ/preview-dataset-human-breast
- Nanostring CosMx Human liver healthy and cancer data: https://nanostring.com/products/cosmx-spatial-molecular-imager/ffpe-dataset/human-liver-rna-ffpe-dataset/
- Vizgen MERSCOPE Human breast cancer data:https://info.vizgen.com/ffpe-showcase?submissionGuid=c9a25730-3fe5-444b-bef8-1a74d51ddefb

## Code availability

All analysis code presented in this manuscript can be found at https://github.com/phipsonlab/jazzPanda_paper. The analysis website (https://phipsonlab.github.io/jazzPanda_workflowr/) was created using the workflowr (1.7.1) R package. The GitHub repository associated with the analysis website is at: https://github.com/phipsonlab/jazzPanda_workflowr. The method is implemented in the open-source Bioconductor R package jazzPanda https://bioconductor.org/packages/jazzPanda.

## Author contributions statement

BP conceived the jazzPanda method and supervised the project. XJ developed the method, implemented the R/Bioconductor package, performed all analyses and simulations, and drafted the manuscript. GHP advised on the software implementation. JC annotated the Xenium human breast cancer samples and provided feedback on the analysis pipeline. MLAL provided biological expertise and interpretation. GKS provided advice on the statistical method and software implementation. BP and XJ wrote the manuscript. All authors reviewed and approved the final manuscript.

## Acknowledgments

We thank the WEHI Research Computing Platform for providing support and access to their High-Performance Computing facility.

