## Supplementary Information for "jazzPanda: spatially aware marker gene detection for imaging-based spatial transcriptomics"

### Supplementary file

### Supplementary Methods

#### Quality control and processing

*Xenium.* Cells with no transcript detections were removed. Low-quality transcript detections, including unassigned detections and those with a quality value below 20, were filtered out.

*CosMx.* For the two CosMx datasets, cells labelled "NotDet" were removed. Transcript coordinates were converted from pixels to micrometres by fitting a linear transformation for each field of view (FOV): using two cells per FOV we estimated the slope and intercept, then mapped every transcript accordingly.

*MERSCOPE.* Cells with fewer than 50 or more than 2,500 total detections were removed.

#### Clustering

*Xenium human breast cancer.* The two samples were processed separately and aligned on their shared cell types. For sample 1 we used the provided cell type labels, collapsing subtypes (for example, $CD4^+$/$CD8^+$ T cells and T-cell–tumour into T_Cells, macrophage subclusters into Macrophages, myoepithelial subclusters into Myoepithelial cells, dendritic subclusters into Dendritic cells, and invasive/DCIS tumour subtypes into Tumor). For sample 2 we performed Louvain clustering (resolution 0.5) and annotated the resulting clusters with known marker genes, giving 14 cell types that were grouped into broader categories for alignment with sample 1 (for example, B and Plasma into B_Cells, and T/NK clusters into T_Cells). Cells from the nine shared cell types in each sample were then joined for downstream analysis.

*MERSCOPE human breast cancer.* We applied the Banksy (1.4.0) pipeline to the log-normalised data (*NormalizeData* from the Seurat package). Banksy PCA and UMAP were computed with lambda 0.2. Clustering was run at resolutions 0.5 and 0.8 (*clusterBanksy*) and refined with *connectClusters*; resolution 0.5 was selected for cell type annotation, giving 10 clusters.

*CosMx human healthy liver and CosMx liver cancer.* The provided cell types were used as cluster labels for these two datasets, with several cell types grouped together (for example, antibody-secreting and mature B cells into B cells, $CD3^+$ alpha-beta and gamma-delta T cells into T cells, and inflammatory and non-inflammatory macrophages into Macrophages).

#### Effect of the bin length on marker gene selection

We assessed marker gene consistency across bin lengths from $10 \times 10$ to $100 \times 100$ for the one-sample CosMx human liver cancer and the two-sample Xenium human breast cancer data.

*Correlation approach.* We used the *compute_permp* function at each bin length. A gene was called a marker when it had a significant adjusted p-value and an observed correlation greater than 0.05, and marker genes were compared across bin lengths for each cluster.

*Linear modelling approach.* For each bin length, spatial vectors were created for every cluster and gene (*get_vectors*) and for the negative controls (*create_genesets*). Each gene was assigned to its most relevant cluster, and the results were summarised for visualisation.

#### Time and memory usage

All jobs were run on the WEHI HPC cluster (SLURM scheduler), each allocated 12 cores (Intel Xeon CPU) and 200–400 GB of memory. SLURM job arrays were used for the simulation and the computational complexity experiments. Runtime and peak memory were measured with the *peakRAM* R package (1.0.2).

#### Computational complexity relative to the number of cells and transcripts

To assess how the time to compute spatial vectors scales with the number of detections, we incrementally generated random points between $10^6$ and $10^9$ (Figure S17). Each transcript count was tested five times, and the runtime and memory for building spatial vectors were measured with *peakRAM* (1.0.2).

#### Computational complexity relative to bin shape and length

We measured the runtime and peak memory for computing 1,000 gene vectors across a range of bin lengths: square bins from $10^2$ ($10 \times 10$) to $10^4$ ($100 \times 100$), and hexagonal bins with side lengths 620, 310, 207, 155, 124, 103, 89 and 78. Both spatial vector creation and linear model fitting were profiled with *peakRAM* (1.0.2), and each bin length was measured five times.

#### Non-specific binding of RNA molecules

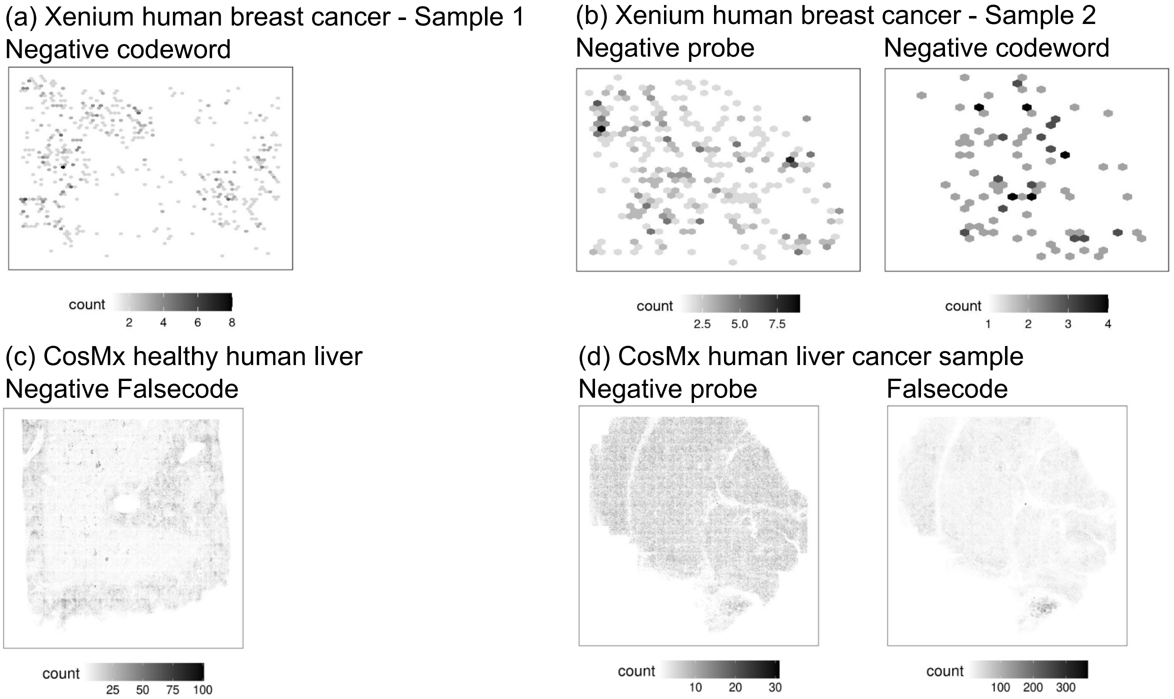

**Figure S1:** Spatial distribution of various negative control signal across multiple datasets: (a) Xenium human breast cancer (Sample 1), negative codeword; (b) Xenium human breast cancer (Sample 2), negative probe and negative codeword; (c) CosMx healthy human liver, negative falsecode; (d) CosMx human liver cancer sample, negative probe and falsecode. Grayscale intensity indicates the number of detected negative control counts.

### Dataset overview

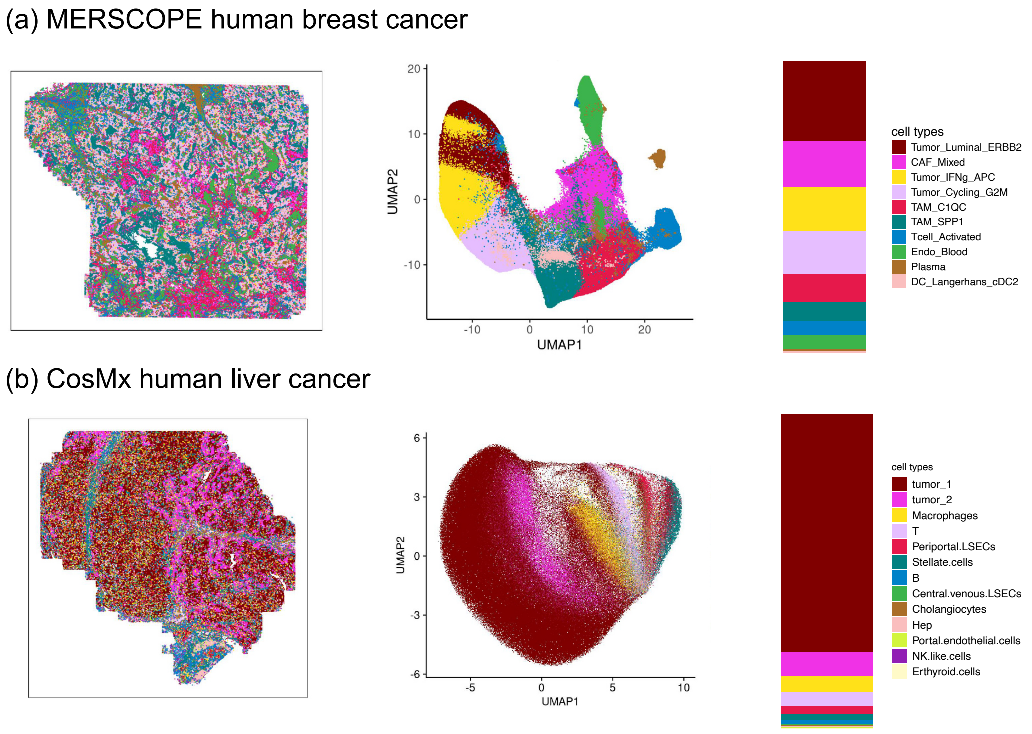

**Figure S2:** Overview of each spatial transcriptomics dataset. Each dataset is represented by three panels: spatial distribution of cell coordinates coloured by annotated cell type, UMAP embedding of the same cells, and cell-type composition shown as relative proportions. Rows correspond to (a) MERSCOPE human breast cancer, and (b) CosMx human liver cancer dataset.

#### Simulation

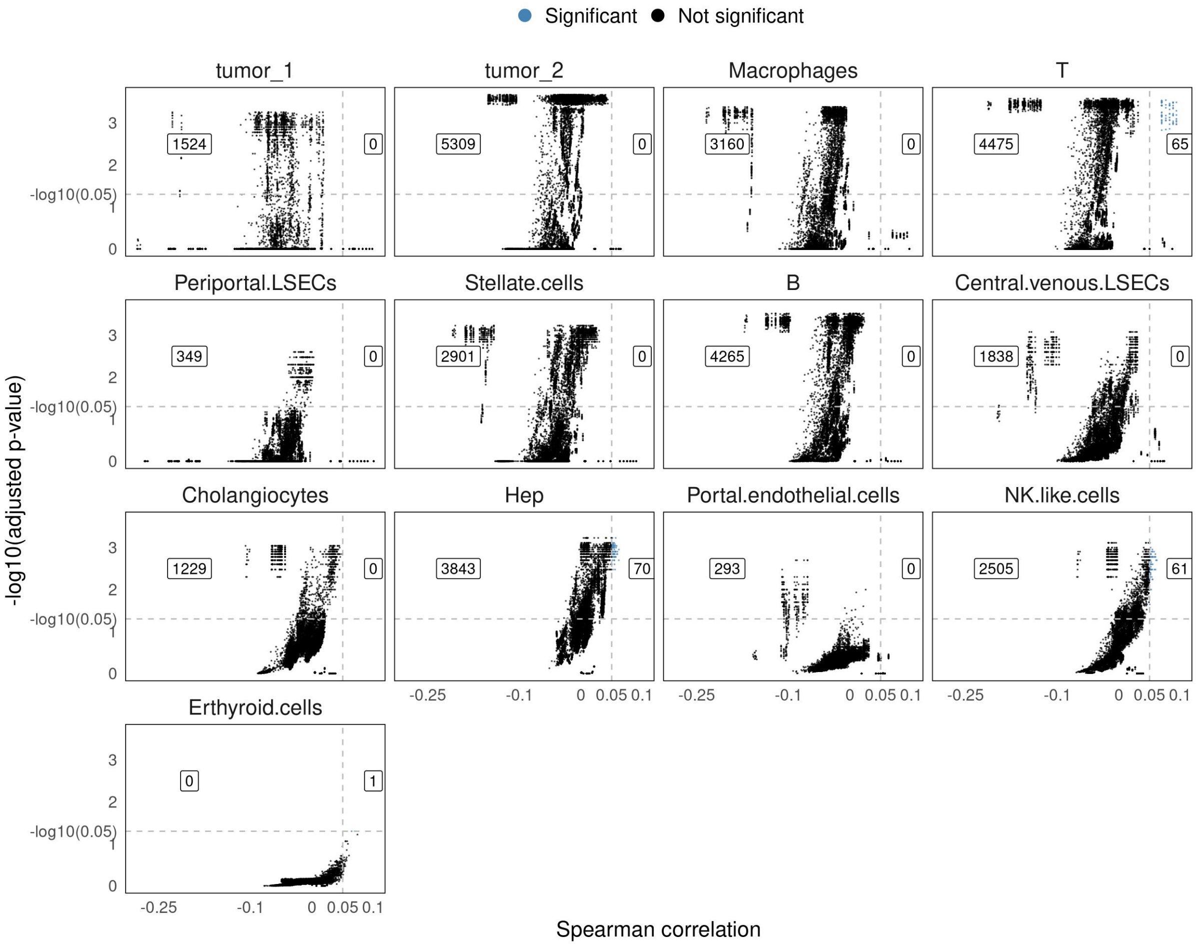

**Figure S3:** Scatter plot showing correlation on x and the adjusted p-values from permutation for each cell type. We show the number of significant genes divided into two numbers for each cell type: the number on the left represents the number of significant genes with correlation<=0.05 while the number on the right refers to the number of significant genes with correlation>0.05

### CosMx healthy human liver (single sample)

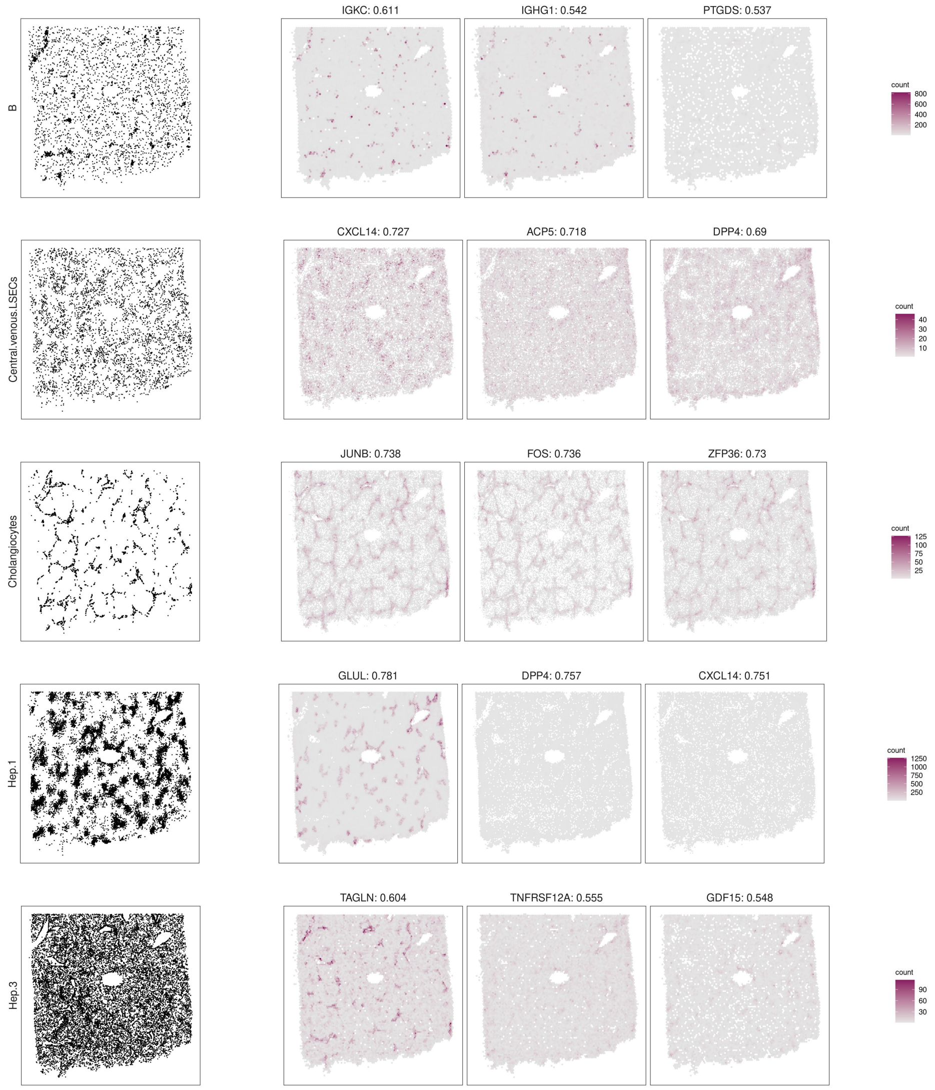

**Figure S4:** Top marker genes identified by jazzPanda-correlation in the CosMx human healthy liver dataset. For each cluster, spatial distributions of cell locations and transcript coordinates are shown for the top three marker genes, selected by minimum adjusted permutation p-value and maximum Spearman correlation.

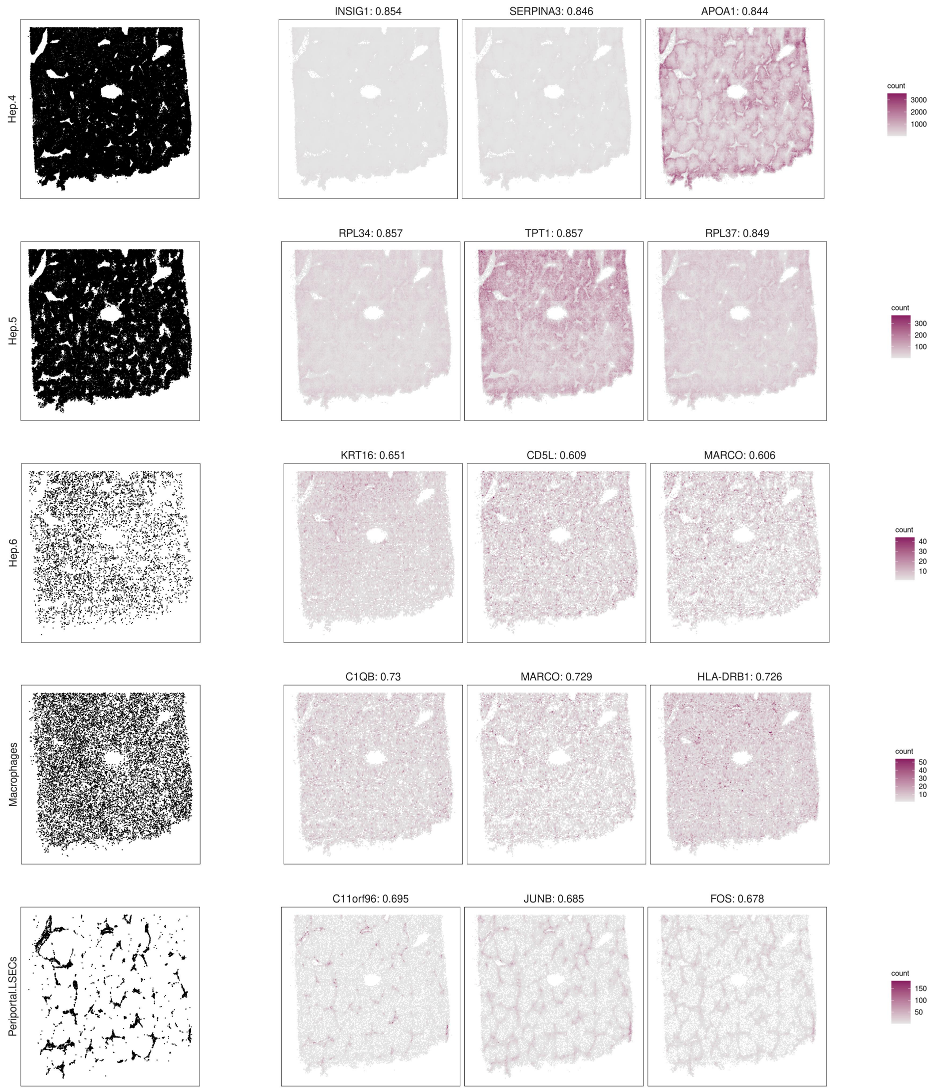

**Figure S4** ***(continued):*** Top marker genes identified by jazzPanda-correlation in the CosMx human healthy liver dataset. For each cluster, spatial distributions of cell locations and transcript coordinates are shown for the top three marker genes, selected by minimum adjusted permutation p-value and maximum Spearman correlation.

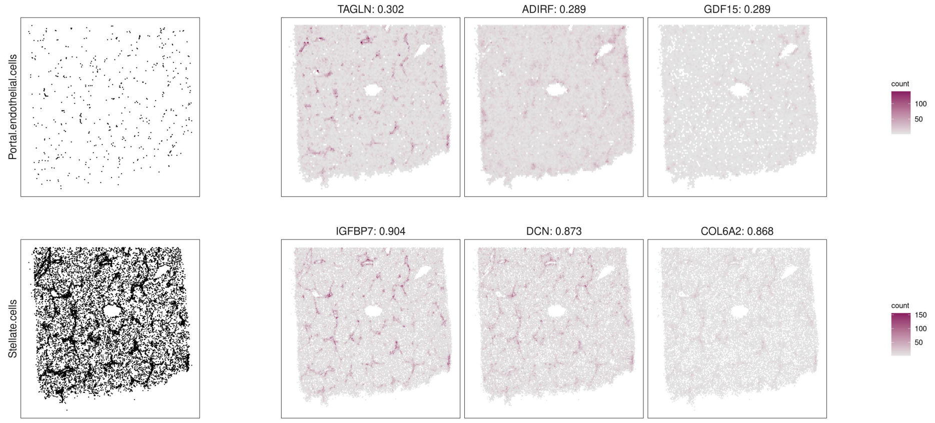

**Figure S4** ***(continued):*** Top marker genes identified by jazzPanda-correlation in the CosMx human healthy liver dataset. For each cluster, spatial distributions of cell locations and transcript coordinates are shown for the top three marker genes, selected by minimum adjusted permutation p-value and maximum Spearman correlation.

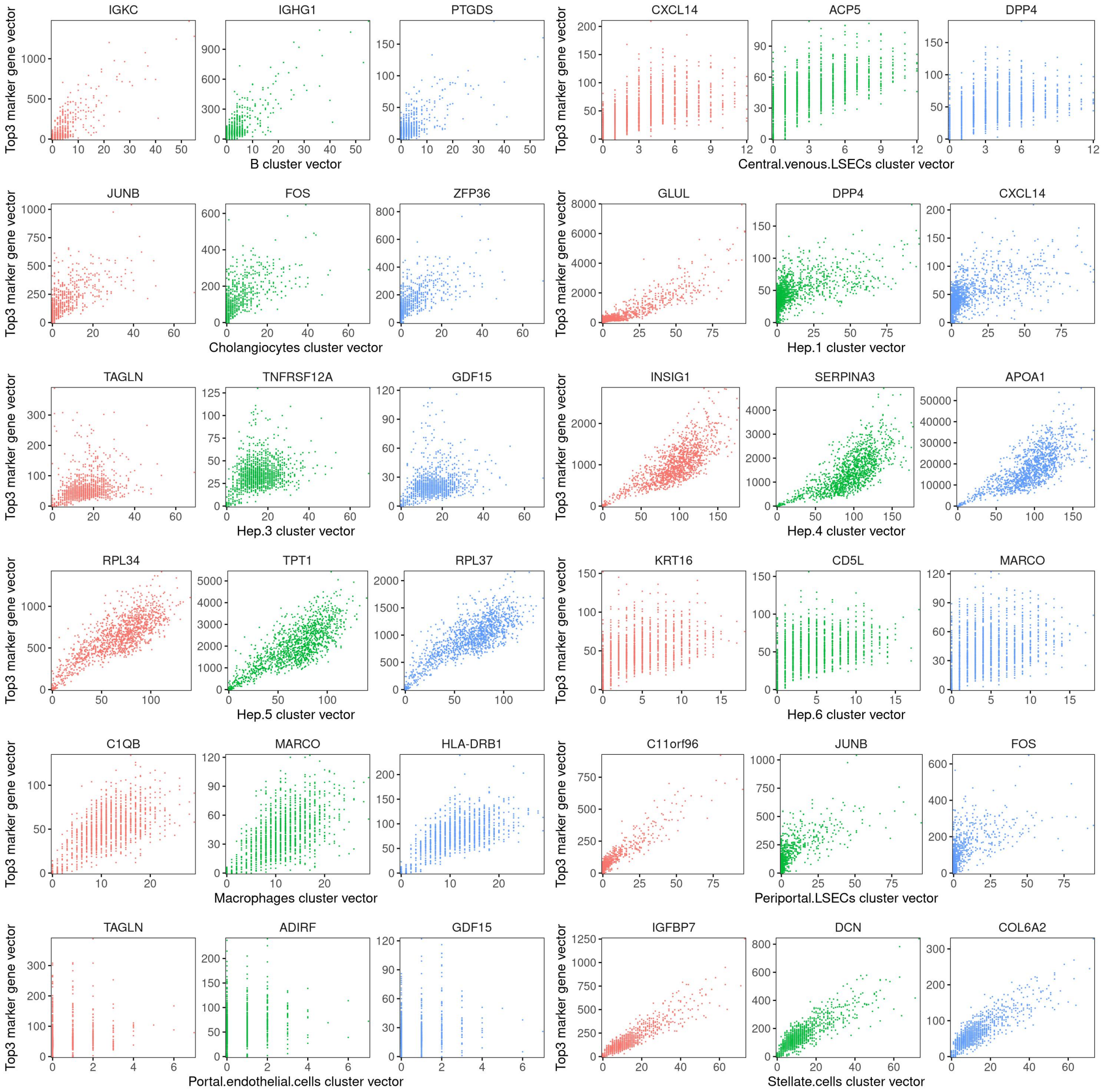

**Figure S5:** Top marker genes identified by jazzPanda-correlation in the CosMx human healthy liver dataset. For each cluster, the relationship between the cluster vector (x-axis) and the top three marker gene vectors (y-axis) is shown. Each point represents a grid bin and is coloured according to the corresponding marker gene identity.

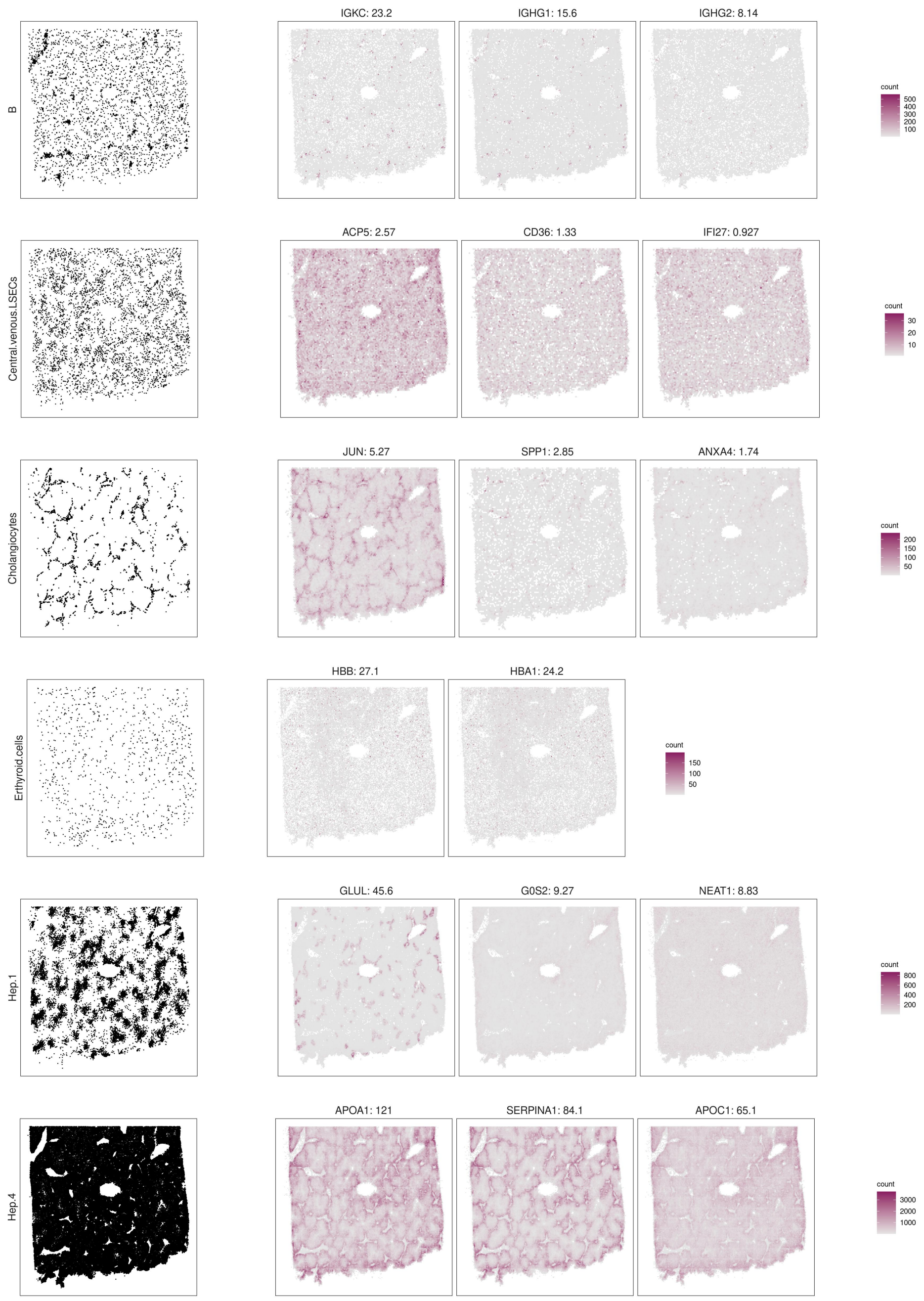

**Figure S6:** Top marker genes identified by jazzPanda-glm in the CosMx human healthy liver dataset. For each cluster, spatial distributions of cell locations and transcript coordinates are shown for the top three marker genes, selected by largest model coefficient.

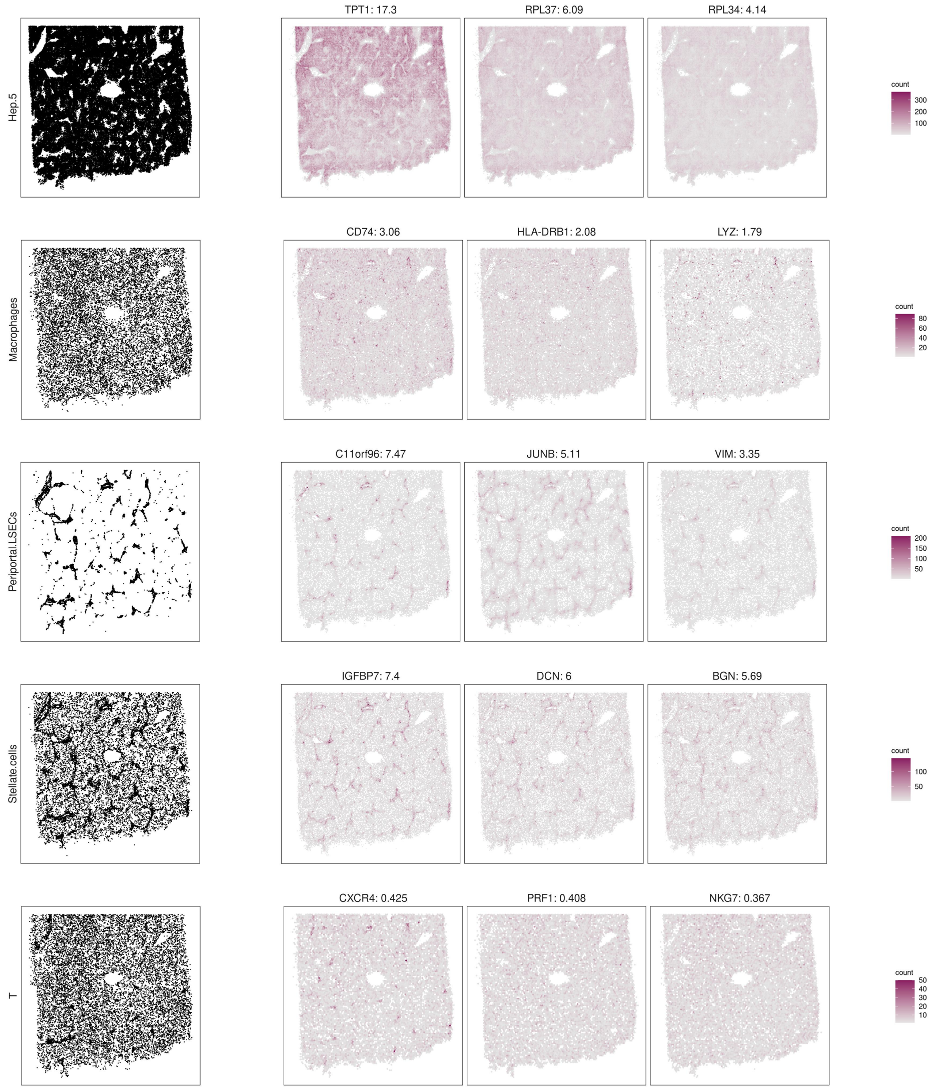

**Figure S6** ***(continued):*** Top marker genes identified by jazzPanda-glm in the CosMx human healthy liver dataset. For each cluster, spatial distributions of cell locations and transcript coordinates are shown for the top three marker genes, selected by largest model coefficient.

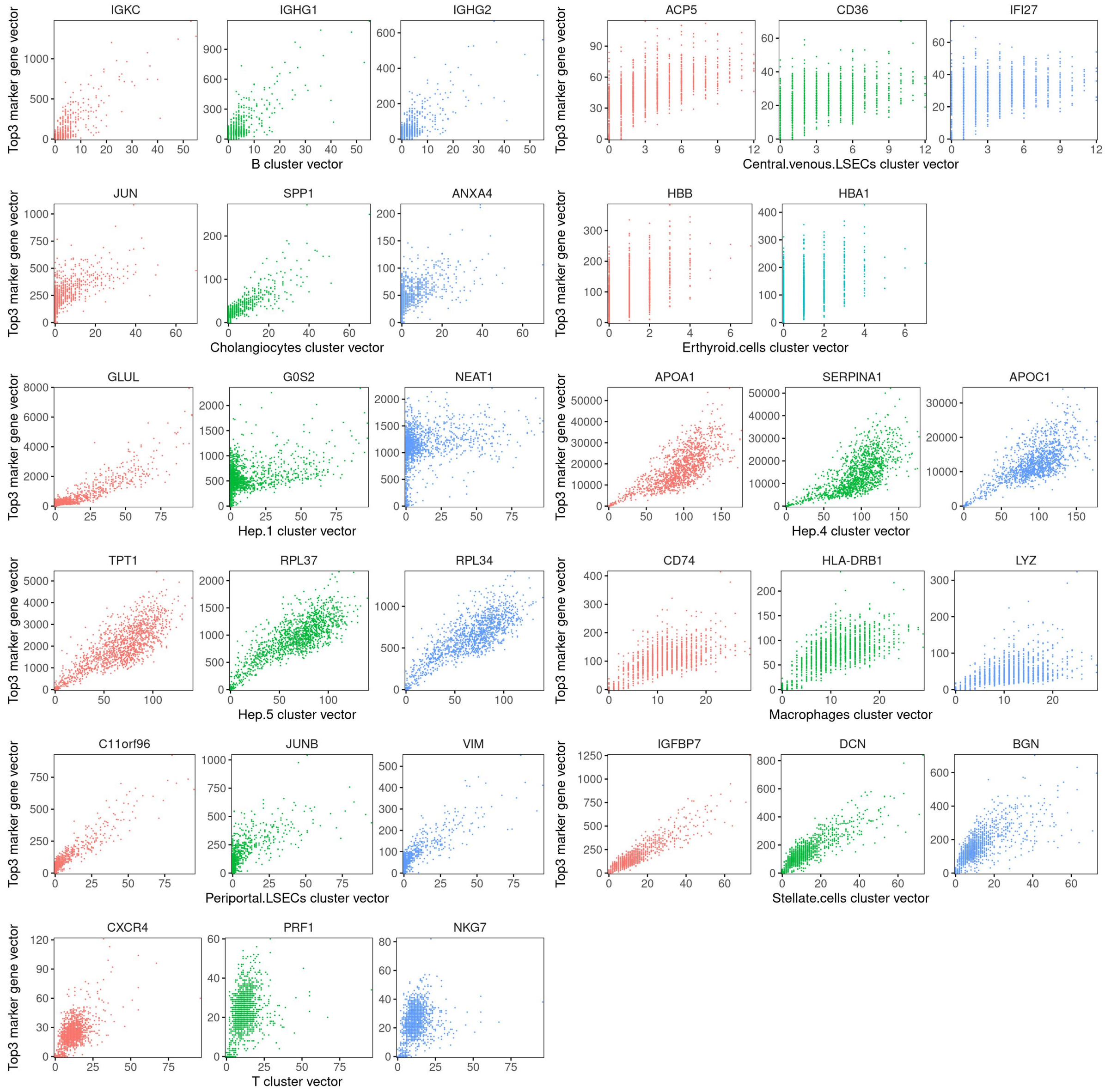

**Figure S7:** Top marker genes identified by jazzPanda-glm in the CosMx human healthy liver dataset. For each cluster, the relationship between the cluster vector (x-axis) and the top three marker gene vectors (y-axis) is shown. Each point represents a grid bin and is coloured according to the corresponding marker gene identity.

### Xenium human breast cancer (multi-sample)

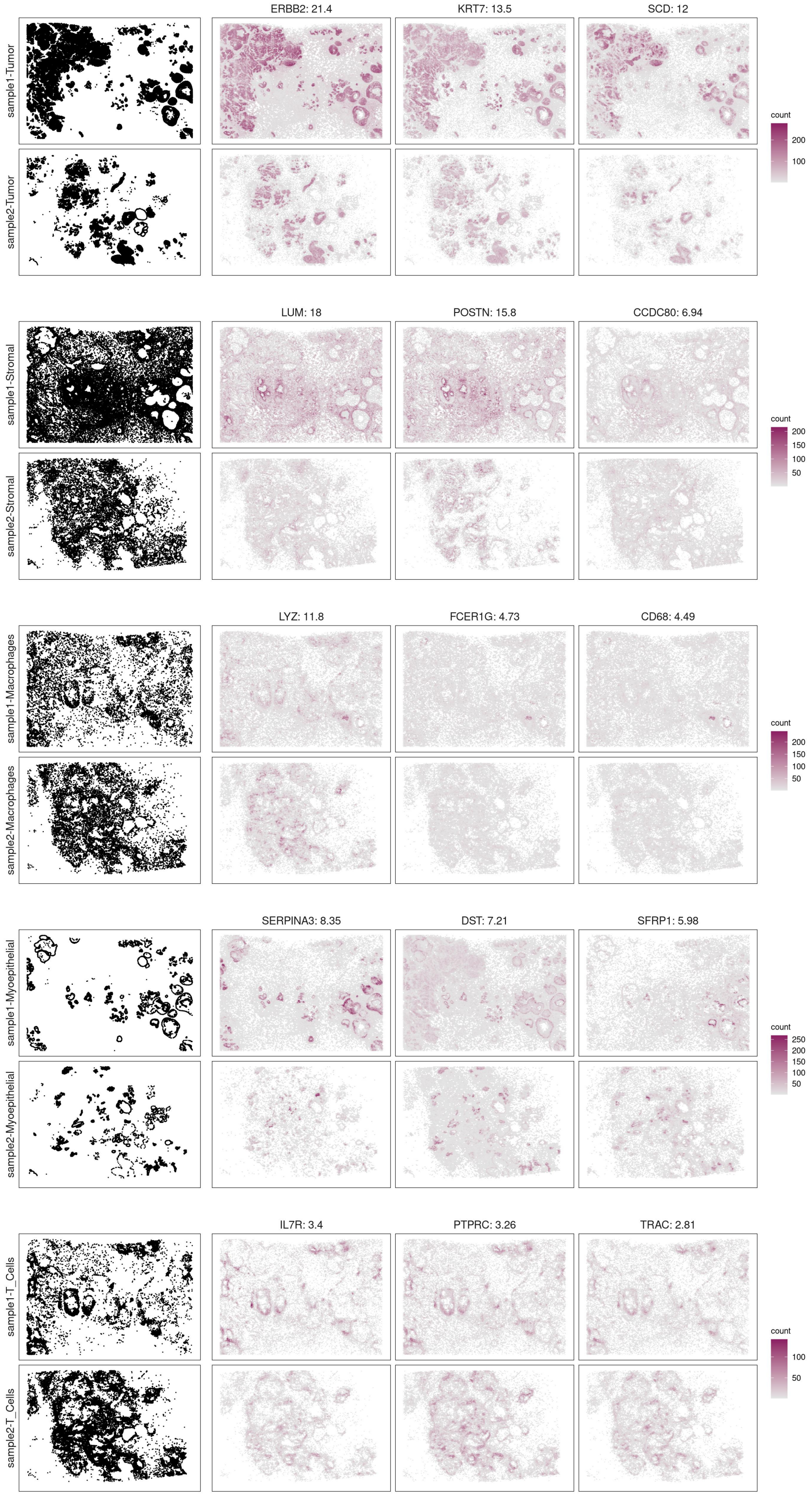

**Figure S8:** Top marker genes identified by jazzPanda-glm in the Xenium human breast cancer dataset. For each cluster, spatial distributions of cell locations and transcript coordinates are shown for the top three marker genes, selected by largest model coefficient.

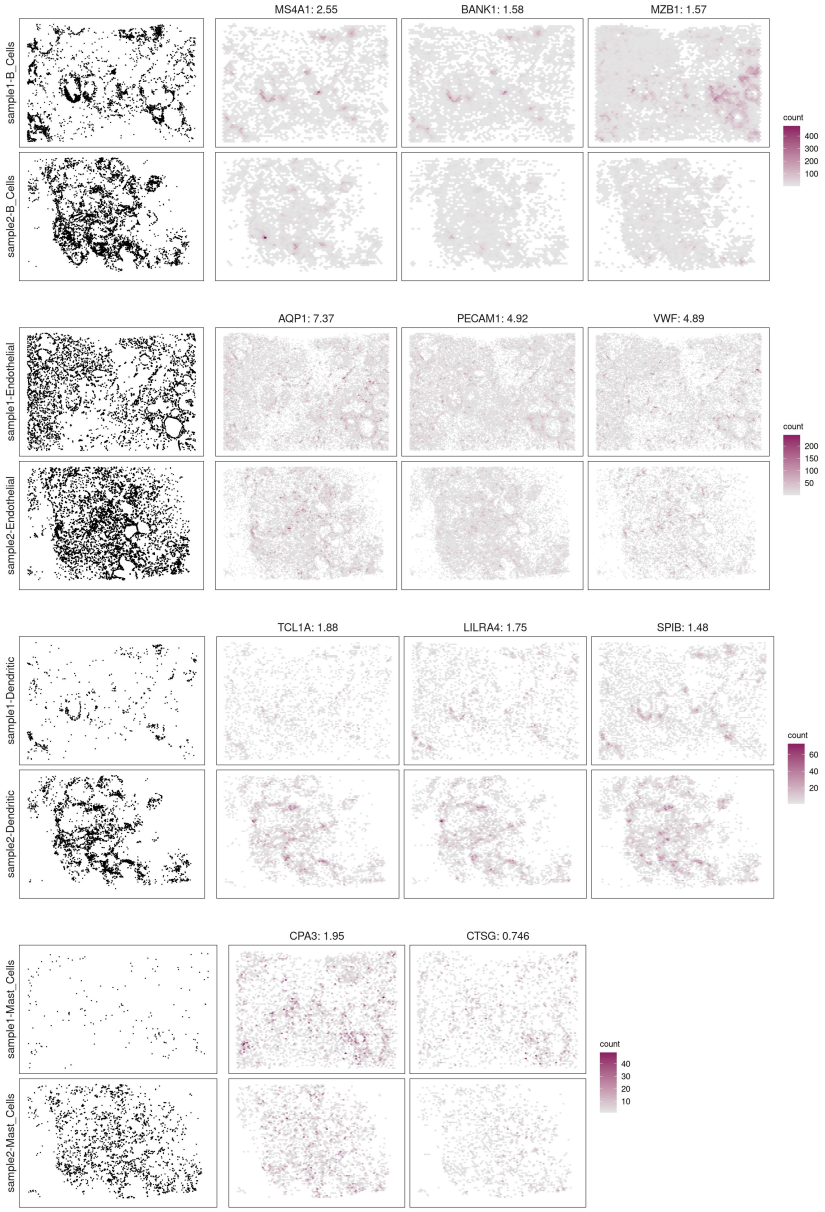

**Figure S8** ***(continued):*** Top marker genes identified by jazzPanda-glm in the Xenium human breast cancer dataset. For each cluster, spatial distributions of cell locations and transcript coordinates are shown for the top three marker genes, selected by largest model coefficient.

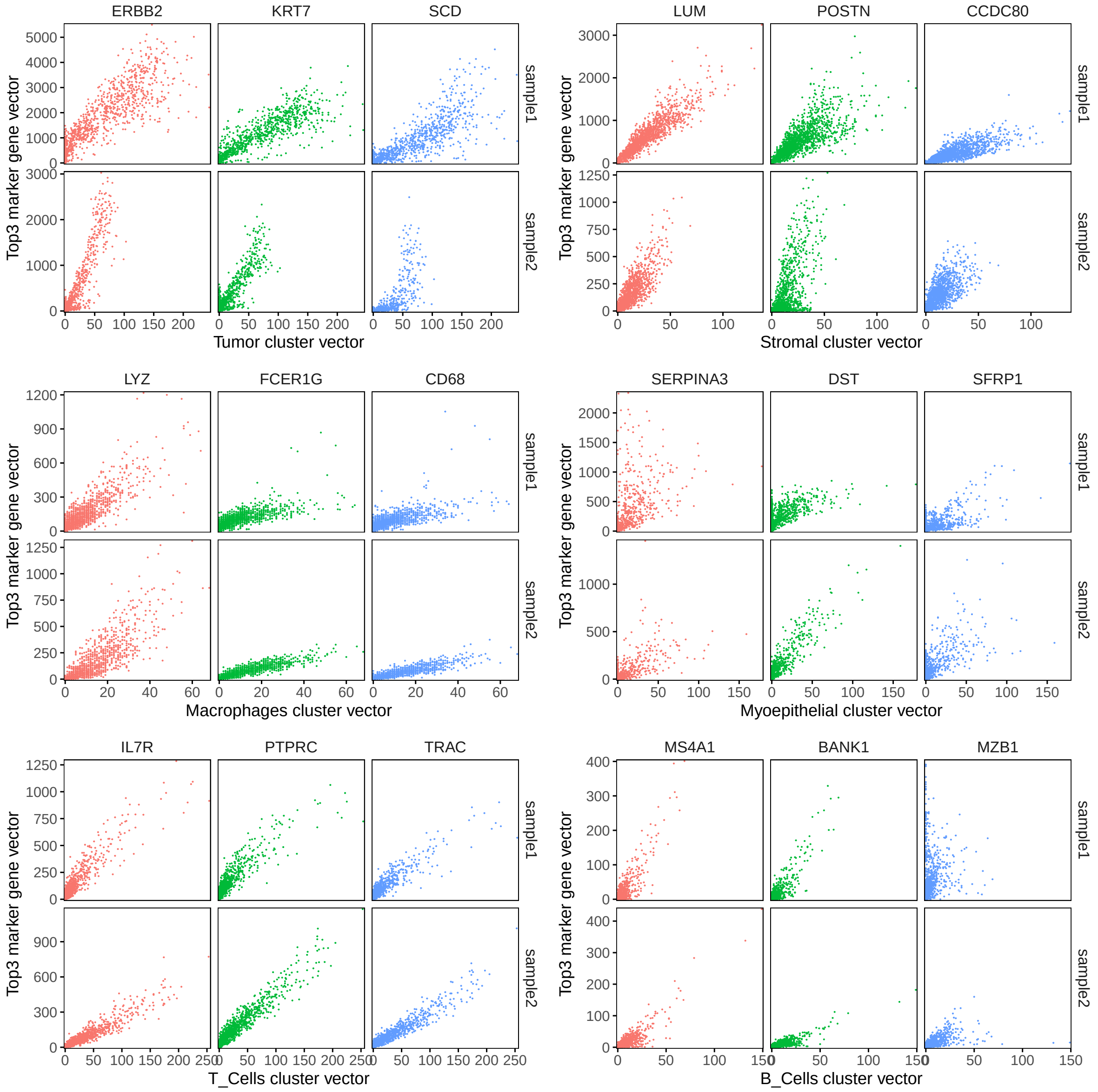
ERBB2
KRT7
SCD
sample1
sample2
Top3 marker gene vector
Tumor cluster vector
LUM
POSTN
CCDC80
Stromal cluster vector
LYZ
FCER1G
CD68
Macrophages cluster vector
SERPINA3
DST
SFRP1
Myoepithelial cluster vector
IL7R
PTPRC
TRAC
T_Cells cluster vector
MS4A1
BANK1
MZB1
B_Cells cluster vector

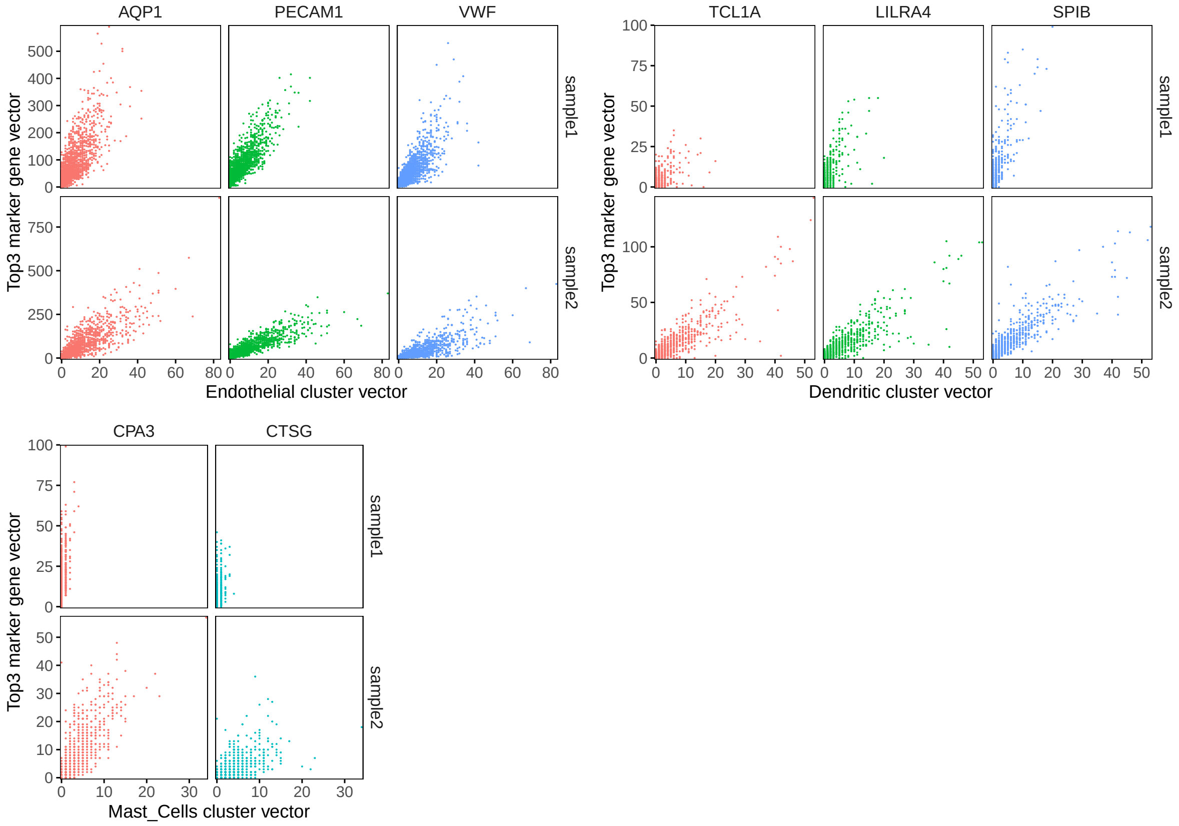

**Figure S9:** Top marker genes identified by jazzPanda-glm in the Xenium human breast cancer dataset. For each cluster, the relationship between the cluster vector (x-axis) and the top three marker gene vectors (y-axis) is shown. Each point represents a grid bin and is coloured according to the corresponding marker gene identity.

### Comparison of markers detected by different methods (upset plots)

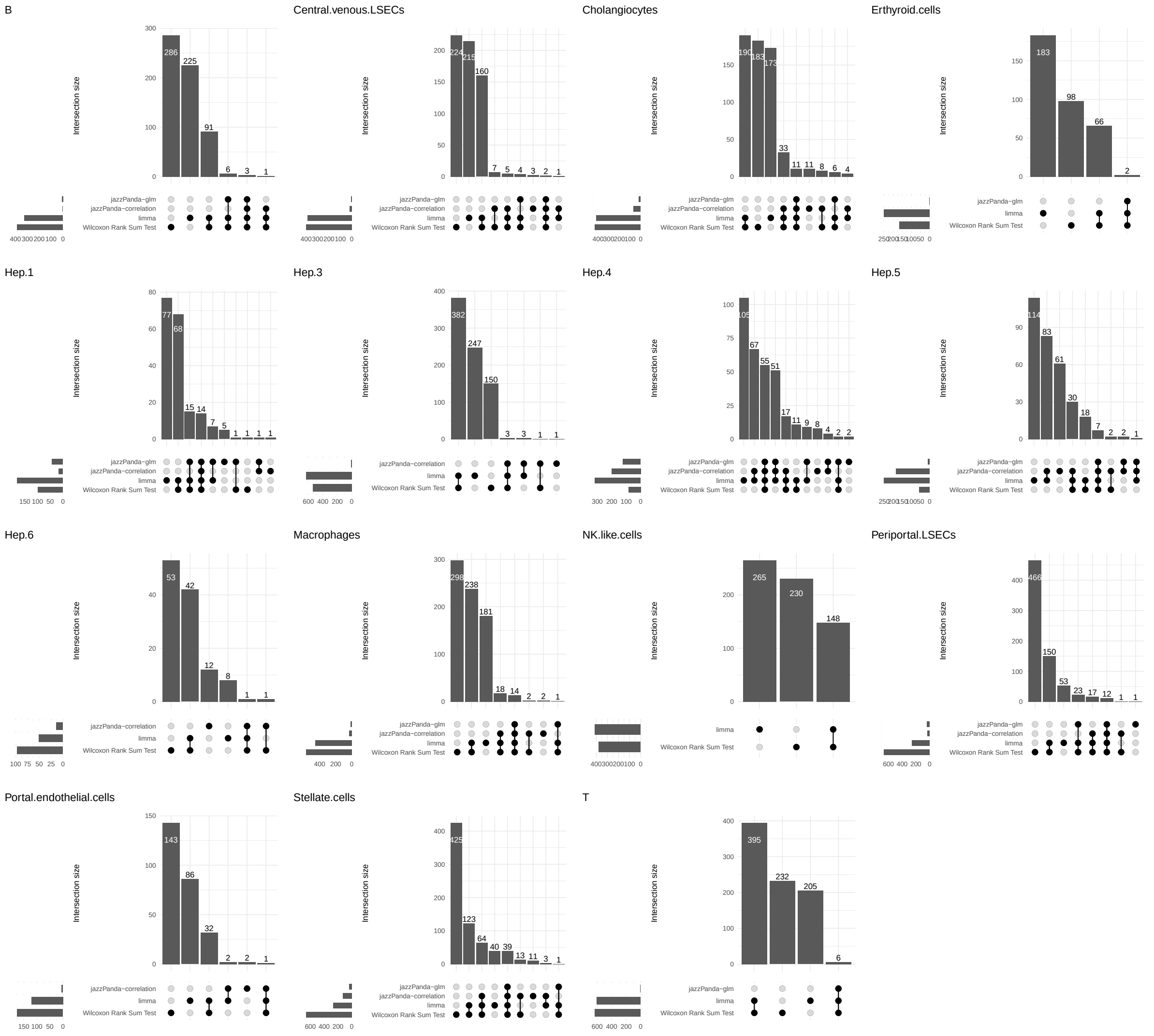

**Figure S10:** Comparison of marker genes identified for the CosMx human healthy liver dataset. For each cluster, marker genes were determined using four approaches: jazzPanda-correlation, jazzPanda-glm, the Wilcoxon Rank Sum Test and limma. The overlap among methods is visualized using an upset plot.

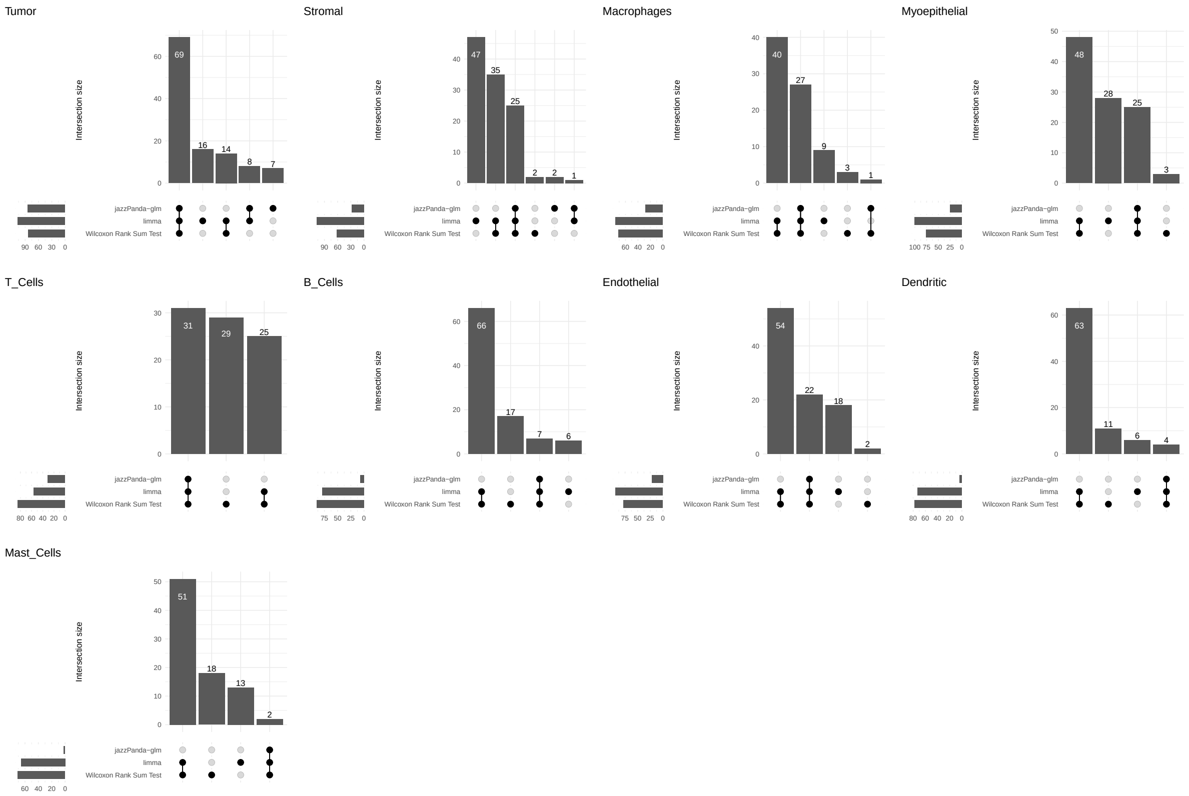

**Figure S11:** Comparison of marker genes identified for the Xenium human breast cancer dataset. For each cluster, marker genes were determined using three approaches: jazzPanda-glm, the Wilcoxon Rank Sum Test and limma. The overlap among methods is visualized using an upset plot.

### Comparison of markers detected by different methods (cumulative correlation)

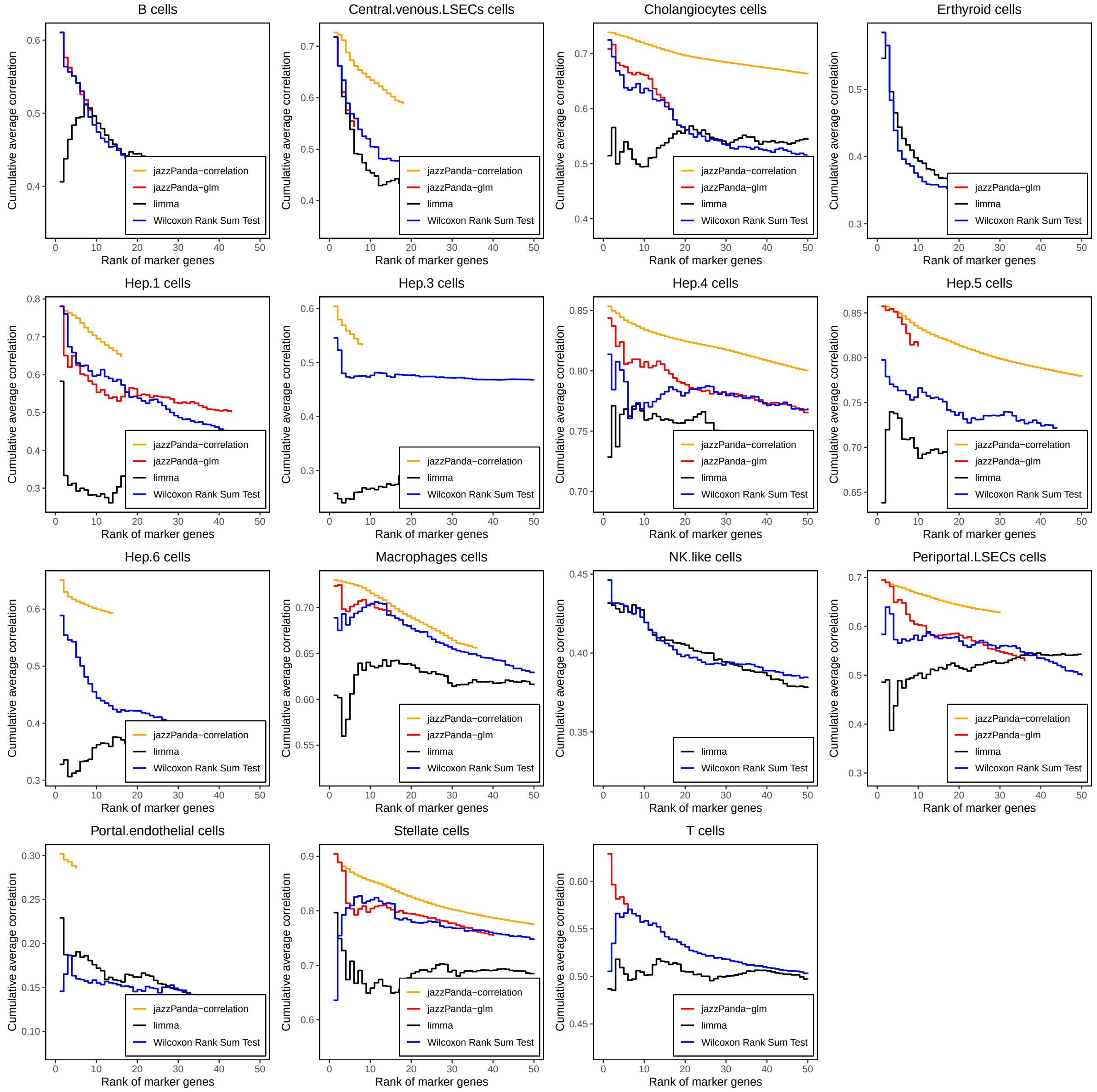

**Figure S12:** Comparison of marker genes identified for the CosMx human healthy liver dataset. For each cluster, marker genes were determined using four approaches: jazzPanda-correlation, jazzPanda-glm, the Wilcoxon Rank Sum Test and limma. The cumulative moving average of the marker–cluster correlation values across the top 50 marker genes is shown for each method.

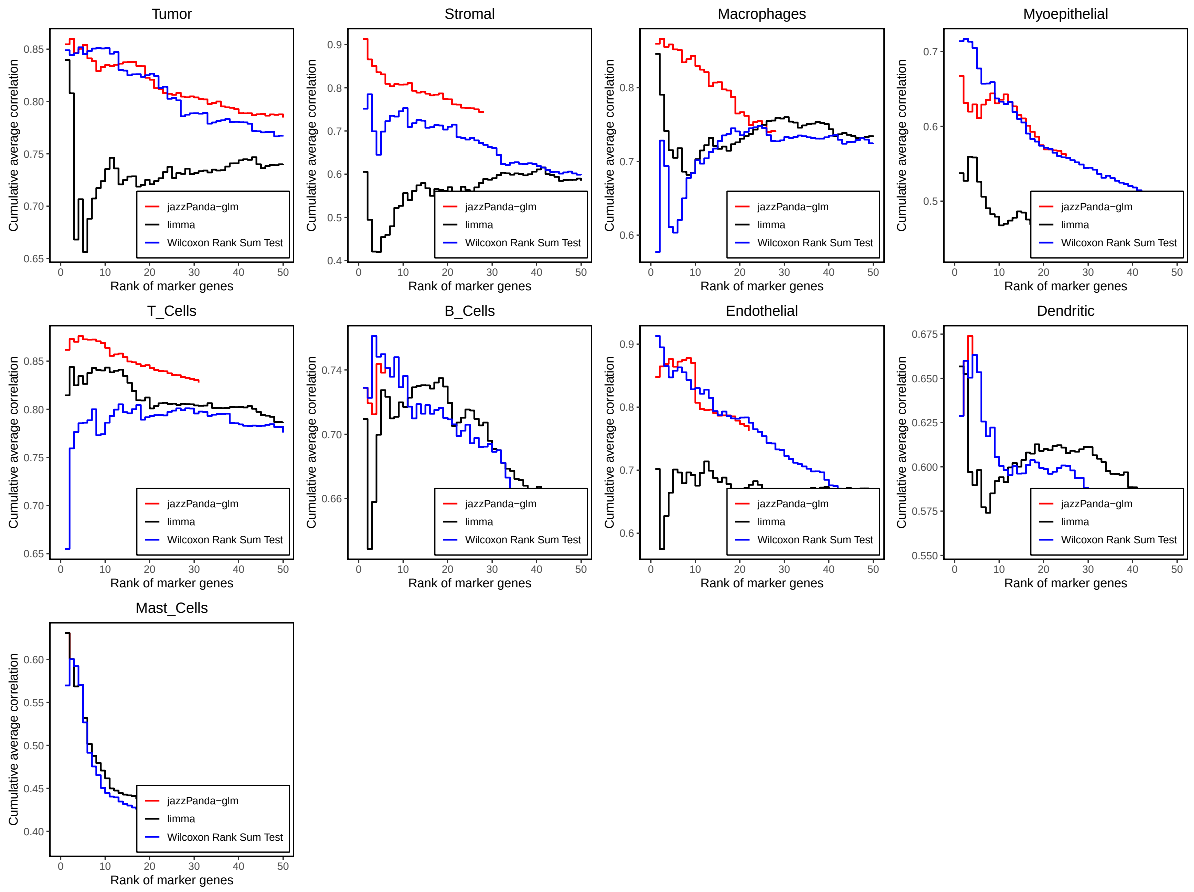

**Figure S13:** Comparison of marker genes identified for the Xenium human breast cancer dataset. For each cluster, marker genes were determined using four approaches: jazzPanda-correlation, jazzPanda-glm, the Wilcoxon Rank Sum Test and limma. The cumulative moving average of the marker–cluster correlation values across the top 50 marker genes is shown for each method.

#### Extension of statistical framework: spatial co-location

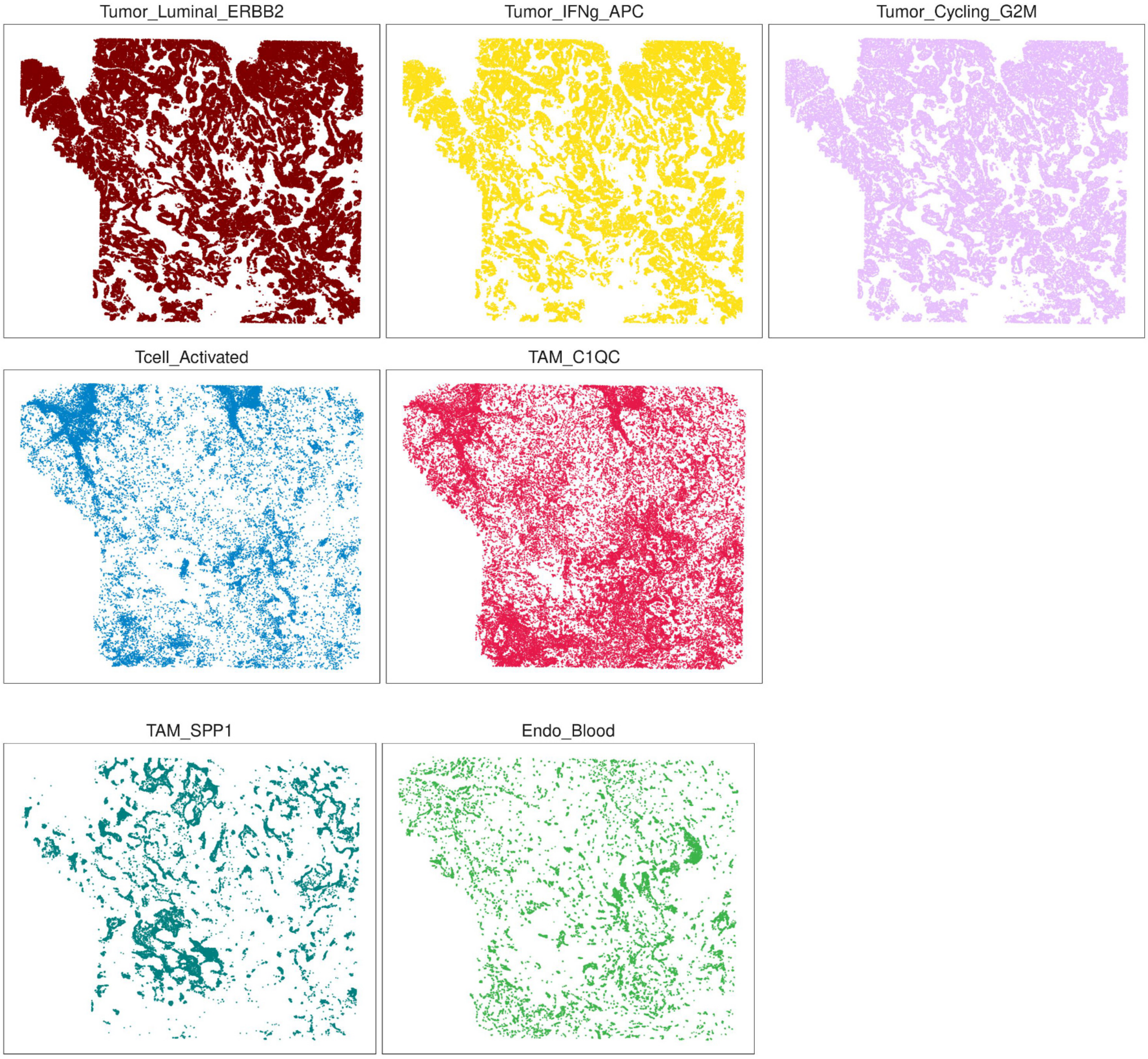

**Figure S14:** Spatial visualisation of Tumor_Luminal_ERBB2, Tumor_IFNg_APC, Tumor_Cycling_G2M,Tcell_Activated, TAM_C1QC, TAM_SPP1 and Endo_Blood in MERSCOPE human breast cancer sample. Each panel shows the tissue coordinates of cells assigned to one cell type, and the points are individual cells.

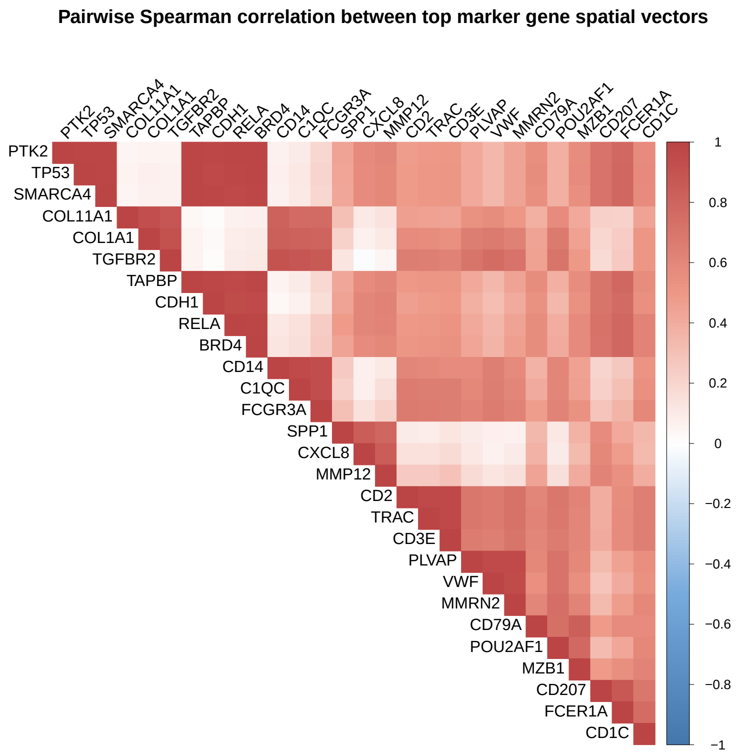

**Figure S15:** Pairwise Spearman correlation between the spatial vectors of top 3 marker genes for each cluster.

### Effect of tile shape and grid length on marker gene detection

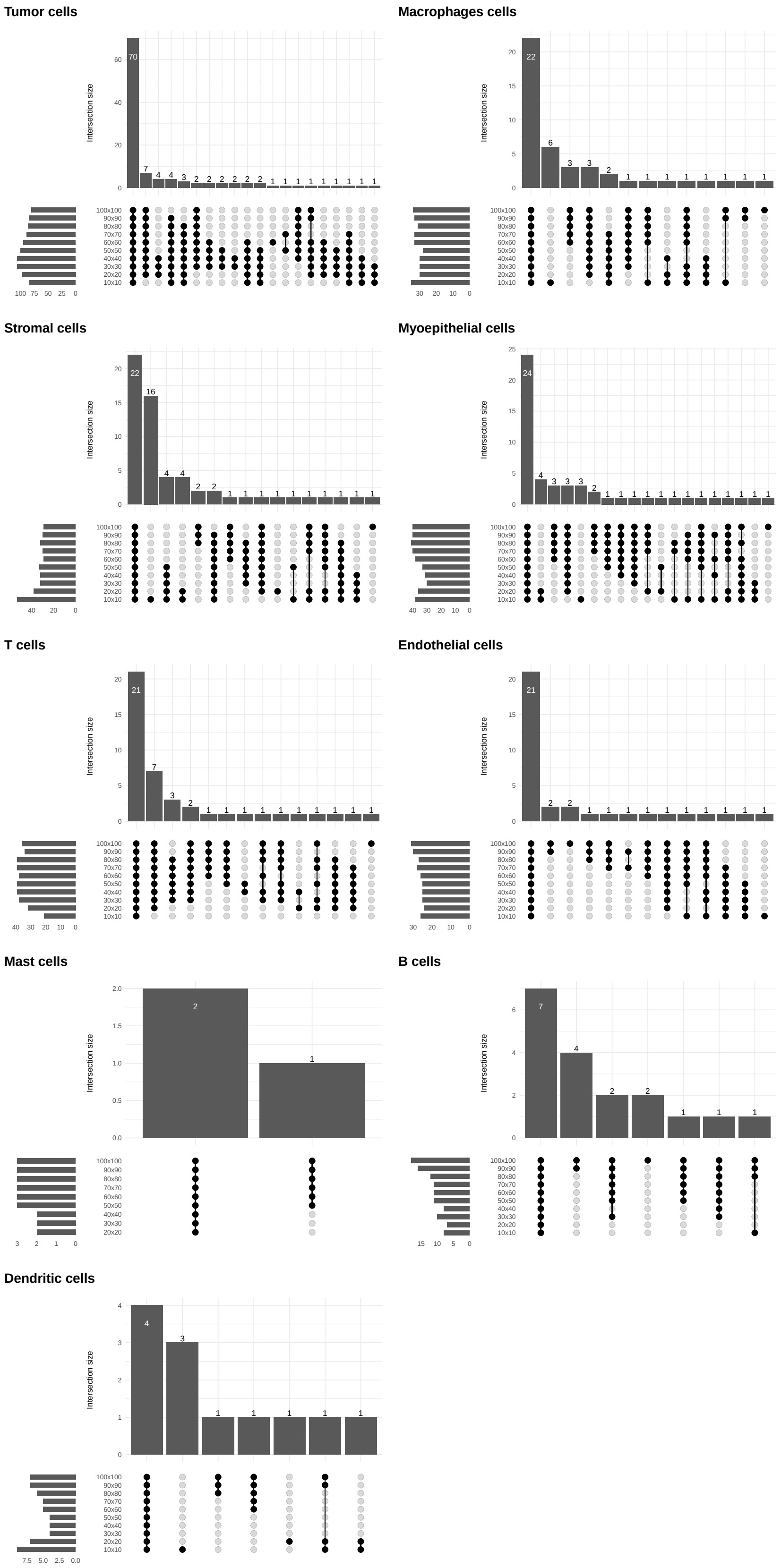

**Figure S16:** A upset diagram illustrating the marker gene result using jazzPanda-glm with different tile length for each cluster (Xenium human breast cancer data).

### Computational complexity

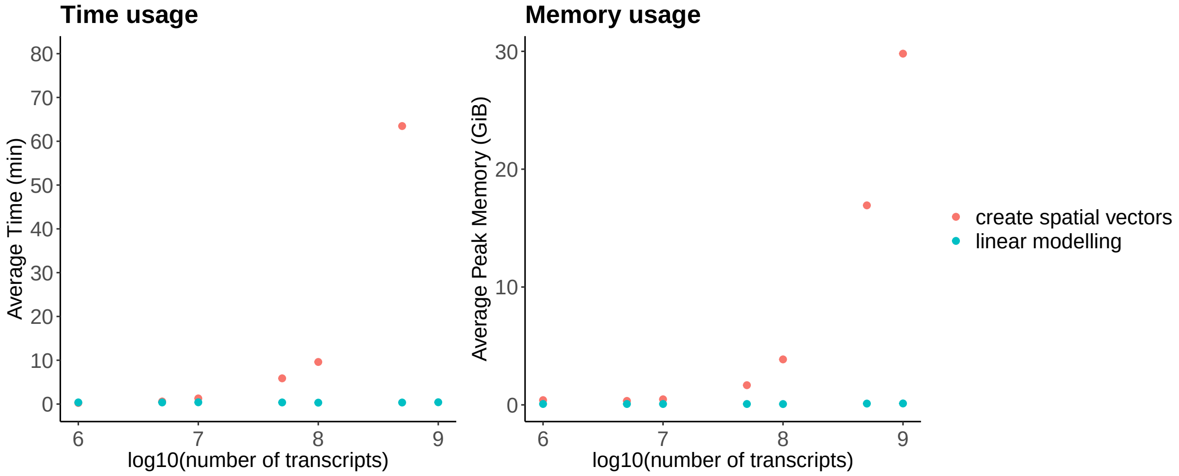

**Figure S17:** The time (left) and memory (right) usage on building spatial vector for different number of genes with 1 core using the CosMx Human Liver cancer data. We incrementally sample between 100 and 1000 genes at an increment of 100 genes per step. Each dot in the figure corresponds to the average of three repeated runs.

(a)

(b)

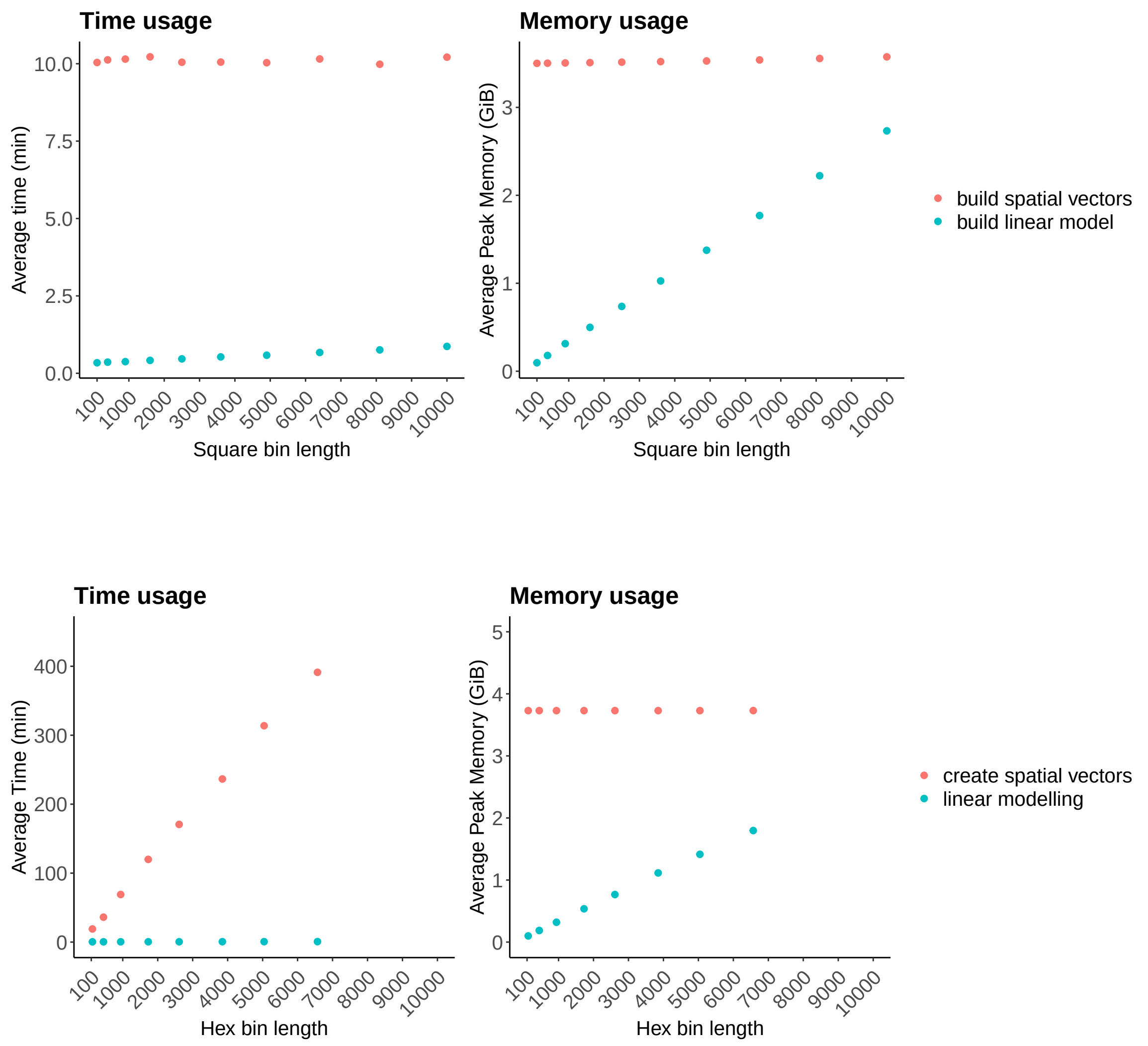

**Figure S18:** Time and memory usage on building spatial vectors and linear modelling for 100 genes across $10^6$ transcripts with varying tile sizes, on a single core. (a) Square bins with the number of tiles ranging from 100 to 10,000. (b) Hex bins with the number of tiles ranging from approximately 100 to 6,300. Each point corresponds to the mean of five repeated runs

### Supplementary Tables

**Table T1:** MERSCOPE human breast cancer data: Spearman correlation matrix between top3 marker genes identified by jazzPanda-glm

| | PTK2 | TP53 | SMARCA4 | COL11A1 | COL1A1 | TGFBR2 | TAPBP | CDH1 | RELA | BRD4 | CD14 | C1QC |
|---|---|---|---|---|---|---|---|---|---|---|---|---|
| PTK2 | 1.00 | 0.99 | 0.99 | 0.04 | 0.06 | 0.06 | 0.99 | 0.97 | 0.98 | 0.99 | 0.07 | 0.10 |
| TP53 | 0.99 | 1.00 | 0.99 | 0.06 | 0.07 | 0.07 | 0.98 | 0.97 | 0.97 | 0.98 | 0.08 | 0.11 |
| SMARCA4 | 0.99 | 0.99 | 1.00 | 0.05 | 0.07 | 0.06 | 0.99 | 0.97 | 0.96 | 0.98 | 0.07 | 0.10 |
| COL11A1 | 0.04 | 0.06 | 0.05 | 1.00 | 0.93 | 0.88 | 0.03 | 0.01 | 0.07 | 0.07 | 0.81 | 0.76 |
| COL1A1 | 0.06 | 0.07 | 0.07 | 0.93 | 1.00 | 0.92 | 0.06 | 0.03 | 0.08 | 0.09 | 0.83 | 0.82 |
| TGFBR2 | 0.06 | 0.07 | 0.06 | 0.88 | 0.92 | 1.00 | 0.05 | 0.02 | 0.09 | 0.10 | 0.91 | 0.89 |
| TAPBP | 0.99 | 0.98 | 0.99 | 0.03 | 0.06 | 0.05 | 1.00 | 0.97 | 0.97 | 0.97 | 0.06 | 0.09 |
| CDH1 | 0.97 | 0.97 | 0.97 | 0.01 | 0.03 | 0.02 | 0.97 | 1.00 | 0.95 | 0.96 | 0.04 | 0.07 |
| RELA | 0.98 | 0.97 | 0.96 | 0.07 | 0.08 | 0.09 | 0.97 | 0.95 | 1.00 | 0.99 | 0.12 | 0.14 |
| BRD4 | 0.99 | 0.98 | 0.98 | 0.07 | 0.09 | 0.10 | 0.97 | 0.96 | 0.99 | 1.00 | 0.12 | 0.15 |
| CD14 | 0.07 | 0.08 | 0.07 | 0.81 | 0.83 | 0.91 | 0.06 | 0.04 | 0.12 | 0.12 | 1.00 | 0.95 |
| C1QC | 0.10 | 0.11 | 0.10 | 0.76 | 0.82 | 0.89 | 0.09 | 0.07 | 0.14 | 0.15 | 0.95 | 1.00 |
| FCGR3A | 0.20 | 0.21 | 0.20 | 0.77 | 0.81 | 0.87 | 0.19 | 0.16 | 0.25 | 0.25 | 0.94 | 0.94 |
| SPP1 | 0.45 | 0.44 | 0.43 | 0.30 | 0.21 | 0.13 | 0.43 | 0.45 | 0.48 | 0.45 | 0.25 | 0.22 |
| CXCL8 | 0.59 | 0.57 | 0.57 | 0.11 | 0.06 | -0.01 | 0.57 | 0.60 | 0.61 | 0.57 | 0.07 | 0.07 |
| MMP12 | 0.60 | 0.59 | 0.59 | 0.14 | 0.11 | 0.04 | 0.60 | 0.62 | 0.62 | 0.59 | 0.11 | 0.15 |
| CD2 | 0.48 | 0.49 | 0.49 | 0.47 | 0.58 | 0.66 | 0.49 | 0.47 | 0.49 | 0.51 | 0.60 | 0.67 |
| TRAC | 0.49 | 0.50 | 0.51 | 0.45 | 0.56 | 0.64 | 0.51 | 0.49 | 0.51 | 0.53 | 0.59 | 0.66 |
| CD3E | 0.50 | 0.51 | 0.51 | 0.44 | 0.55 | 0.62 | 0.52 | 0.50 | 0.52 | 0.54 | 0.58 | 0.65 |
| PLVAP | 0.41 | 0.42 | 0.42 | 0.53 | 0.65 | 0.71 | 0.42 | 0.39 | 0.41 | 0.43 | 0.57 | 0.59 |
| VWF | 0.35 | 0.35 | 0.35 | 0.56 | 0.67 | 0.75 | 0.34 | 0.31 | 0.35 | 0.37 | 0.65 | 0.65 |
| MMRN2 | 0.45 | 0.45 | 0.44 | 0.51 | 0.63 | 0.71 | 0.44 | 0.41 | 0.45 | 0.47 | 0.59 | 0.60 |
| CD79A | 0.55 | 0.56 | 0.56 | 0.38 | 0.44 | 0.46 | 0.57 | 0.53 | 0.57 | 0.57 | 0.38 | 0.40 |
| POU2AF1 | 0.38 | 0.38 | 0.38 | 0.58 | 0.68 | 0.70 | 0.38 | 0.35 | 0.39 | 0.40 | 0.60 | 0.61 |
| MZB1 | 0.57 | 0.57 | 0.56 | 0.42 | 0.46 | 0.50 | 0.57 | 0.53 | 0.59 | 0.59 | 0.46 | 0.46 |
| CD207 | 0.71 | 0.72 | 0.71 | 0.22 | 0.20 | 0.18 | 0.70 | 0.70 | 0.72 | 0.72 | 0.21 | 0.23 |
| FCER1A | 0.78 | 0.79 | 0.78 | 0.22 | 0.25 | 0.25 | 0.78 | 0.77 | 0.78 | 0.79 | 0.27 | 0.31 |
| CD1C | 0.58 | 0.58 | 0.57 | 0.45 | 0.51 | 0.52 | 0.57 | 0.54 | 0.61 | 0.62 | 0.52 | 0.54 |

| | FCGR3A | SPP1 | CXCL8 | MMP12 | CD2 | TRAC | CD3E | PLVAP | VWF | MMRN2 | CD79A | POU2AF1 | MZB1 | CD207 | FCER1A | CD1C |
|---|---|---|---|---|---|---|---|---|---|---|---|---|---|---|---|---|
| PTK2 | 0.20 | 0.45 | 0.59 | 0.60 | 0.48 | 0.49 | 0.50 | 0.41 | 0.35 | 0.45 | 0.55 | 0.38 | 0.57 | 0.71 | 0.78 | 0.58 |
| TP53 | 0.21 | 0.44 | 0.57 | 0.59 | 0.49 | 0.50 | 0.51 | 0.42 | 0.35 | 0.45 | 0.56 | 0.38 | 0.57 | 0.72 | 0.79 | 0.58 |
| SMARCA4 | 0.20 | 0.43 | 0.57 | 0.59 | 0.49 | 0.51 | 0.51 | 0.42 | 0.35 | 0.44 | 0.56 | 0.38 | 0.56 | 0.71 | 0.78 | 0.57 |
| COL11A1 | 0.77 | 0.30 | 0.11 | 0.14 | 0.47 | 0.45 | 0.44 | 0.53 | 0.56 | 0.51 | 0.38 | 0.58 | 0.42 | 0.22 | 0.22 | 0.45 |
| COL1A1 | 0.81 | 0.21 | 0.06 | 0.11 | 0.58 | 0.56 | 0.55 | 0.65 | 0.67 | 0.63 | 0.44 | 0.68 | 0.46 | 0.20 | 0.25 | 0.51 |
| TGFBR2 | 0.87 | 0.13 | -0.01 | 0.04 | 0.66 | 0.64 | 0.62 | 0.71 | 0.75 | 0.71 | 0.46 | 0.70 | 0.50 | 0.18 | 0.25 | 0.52 |
| TAPBP | 0.19 | 0.43 | 0.57 | 0.60 | 0.49 | 0.51 | 0.52 | 0.42 | 0.34 | 0.44 | 0.57 | 0.38 | 0.57 | 0.70 | 0.78 | 0.57 |
| CDH1 | 0.16 | 0.45 | 0.60 | 0.62 | 0.47 | 0.49 | 0.50 | 0.39 | 0.31 | 0.41 | 0.53 | 0.35 | 0.53 | 0.70 | 0.77 | 0.54 |
| RELA | 0.25 | 0.48 | 0.61 | 0.62 | 0.49 | 0.51 | 0.52 | 0.41 | 0.35 | 0.45 | 0.57 | 0.39 | 0.59 | 0.72 | 0.78 | 0.61 |
| BRD4 | 0.25 | 0.45 | 0.57 | 0.59 | 0.51 | 0.53 | 0.54 | 0.43 | 0.37 | 0.47 | 0.57 | 0.40 | 0.59 | 0.72 | 0.79 | 0.62 |
| CD14 | 0.94 | 0.25 | 0.07 | 0.11 | 0.60 | 0.59 | 0.58 | 0.57 | 0.65 | 0.59 | 0.38 | 0.60 | 0.46 | 0.21 | 0.27 | 0.52 |
| C1QC | 0.94 | 0.22 | 0.07 | 0.15 | 0.67 | 0.66 | 0.65 | 0.59 | 0.65 | 0.60 | 0.40 | 0.61 | 0.46 | 0.23 | 0.31 | 0.54 |
| FCGR3A | 1.00 | 0.31 | 0.15 | 0.21 | 0.68 | 0.66 | 0.66 | 0.62 | 0.66 | 0.63 | 0.48 | 0.61 | 0.54 | 0.28 | 0.37 | 0.59 |
| SPP1 | 0.31 | 1.00 | 0.85 | 0.80 | 0.09 | 0.09 | 0.12 | 0.10 | 0.08 | 0.06 | 0.35 | 0.12 | 0.37 | 0.58 | 0.41 | 0.35 |
| CXCL8 | 0.15 | 0.85 | 1.00 | 0.84 | 0.14 | 0.15 | 0.17 | 0.12 | 0.07 | 0.09 | 0.37 | 0.09 | 0.34 | 0.59 | 0.48 | 0.33 |
| MMP12 | 0.21 | 0.80 | 0.84 | 1.00 | 0.25 | 0.26 | 0.30 | 0.18 | 0.12 | 0.16 | 0.44 | 0.15 | 0.41 | 0.62 | 0.55 | 0.39 |
| CD2 | 0.68 | 0.09 | 0.14 | 0.25 | 1.00 | 0.96 | 0.96 | 0.69 | 0.67 | 0.73 | 0.60 | 0.69 | 0.61 | 0.40 | 0.57 | 0.65 |
| TRAC | 0.66 | 0.09 | 0.15 | 0.26 | 0.96 | 1.00 | 0.97 | 0.69 | 0.67 | 0.72 | 0.61 | 0.69 | 0.61 | 0.39 | 0.58 | 0.64 |
| CD3E | 0.66 | 0.12 | 0.17 | 0.30 | 0.96 | 0.97 | 1.00 | 0.67 | 0.65 | 0.70 | 0.61 | 0.67 | 0.61 | 0.41 | 0.58 | 0.65 |
| PLVAP | 0.62 | 0.10 | 0.12 | 0.18 | 0.69 | 0.69 | 0.67 | 1.00 | 0.95 | 0.96 | 0.60 | 0.72 | 0.59 | 0.33 | 0.45 | 0.55 |
| VWF | 0.66 | 0.08 | 0.07 | 0.12 | 0.67 | 0.67 | 0.65 | 0.95 | 1.00 | 0.94 | 0.55 | 0.72 | 0.56 | 0.29 | 0.40 | 0.54 |
| MMRN2 | 0.63 | 0.06 | 0.09 | 0.16 | 0.73 | 0.72 | 0.70 | 0.96 | 0.94 | 1.00 | 0.60 | 0.74 | 0.61 | 0.34 | 0.47 | 0.59 |
| CD79A | 0.48 | 0.35 | 0.37 | 0.44 | 0.60 | 0.61 | 0.61 | 0.60 | 0.55 | 0.60 | 1.00 | 0.73 | 0.84 | 0.48 | 0.57 | 0.58 |
| POU2AF1 | 0.61 | 0.12 | 0.09 | 0.15 | 0.69 | 0.69 | 0.67 | 0.72 | 0.72 | 0.74 | 0.73 | 1.00 | 0.78 | 0.33 | 0.43 | 0.60 |
| MZB1 | 0.54 | 0.37 | 0.34 | 0.41 | 0.61 | 0.61 | 0.61 | 0.59 | 0.56 | 0.61 | 0.84 | 0.78 | 1.00 | 0.48 | 0.56 | 0.61 |
| CD207 | 0.28 | 0.58 | 0.59 | 0.62 | 0.40 | 0.39 | 0.41 | 0.33 | 0.29 | 0.34 | 0.48 | 0.33 | 0.48 | 1.00 | 0.88 | 0.69 |
| FCER1A | 0.37 | 0.41 | 0.48 | 0.55 | 0.57 | 0.58 | 0.58 | 0.45 | 0.40 | 0.47 | 0.57 | 0.43 | 0.56 | 0.88 | 1.00 | 0.75 |
| CD1C | 0.59 | 0.35 | 0.33 | 0.39 | 0.65 | 0.64 | 0.65 | 0.55 | 0.54 | 0.59 | 0.58 | 0.60 | 0.61 | 0.69 | 0.75 | 1.00 |

**Table T2:** Xenium human breast cancer data: Pairwise Jaccard index of detected marker genes for each cluster across different binning resolutions (10×10–100×100). Shared = genes shared by all ten bins; Union = genes in at least one; Mean Jaccard = mean pairwise Jaccard index over the 45 bin pairs

| cluster | cell type | # Shared | # Union | Mean Jaccard |
|---|---|---|---|---|
| c1 | Tumor | 70 | 109 | 0.8540 |
| c3 | Macrophages | 22 | 44 | 0.7940 |
| c2 | Stromal | 22 | 60 | 0.7270 |
| c4 | Myoepithelial | 24 | 52 | 0.7440 |
| c5 | T | 21 | 43 | 0.7890 |
| c7 | Endothelial | 21 | 35 | 0.8270 |
| c9 | Mast | 0 | 3 | 0.6670 |
| c6 | B | 7 | 18 | 0.7070 |
| c8 | Dendritic | 4 | 12 | 0.6610 |
